# Transition from Transient DNA Rereplication to Inherited Gene Amplification Following Prolonged Environmental Stress

**DOI:** 10.1101/2023.05.08.539886

**Authors:** Gregory M Wright, Johannes Menzel, Philip D. Tatman, Joshua C. Black

**Affiliations:** University of Colorado Anschutz Medical Campus, Department of Pharmacology, 12800 E 19^th^ Ave, Aurora CO 80045, USA

**Author notes:** Correspondence should be addressed to: Joshua C. Black Ph.D. University of Colorado School of Medicine Anschutz Medical Campus Mail Stop 8303 12800 E 19^th^ Ave, Aurora CO 80045.

## Abstract

Cells require the ability to adapt to changing environmental conditions, however, it is unclear how these changes elicit stable permanent changes in genomes. We demonstrate that, in response to environmental metal exposure, the metallothionein (MT) locus undergoes DNA rereplication generating transient site-specific gene amplifications (TSSGs). Chronic metal exposure allows transition from MT TSSG to inherited MT gene amplification through homologous recombination within and outside of the MT locus. DNA rereplication of the MT locus is suppressed by H3K27me3 and EZH2. Long-term ablation of EZH2 activity eventually leads to integration and inheritance of MT gene amplifications without the selective pressure of metal exposure. The rereplication and inheritance of MT gene amplification is an evolutionarily conserved response to environmental metal from yeast to human. Our results describe a new paradigm for adaptation to environmental stress where targeted, transient DNA rereplication precedes stable inherited gene amplification.

## INTRODUCTION

Cells and organisms are exposed to constantly changing environmental conditions, which necessitate adaptation though changes in gene expression programs ^1,2^. Significant work has explored how acute changes in the environment alter gene expression ^3–5^, however, it remains unclear how cells and organisms adapt to a chronic or permanently changed environment.

Genomic plasticity has emerged as a key mechanism for cellular adaptation ^6–9^. Changes in cellular genetics including amplifications, deletions, mutations and structural variants have all been linked to changes in cellular environment ^8^. It remains an open question if these events occur at random and are then selected for, or if cellular programs identify and regulate regions of increased genomic plasticity.

Environmentally induced gene amplifications have been reported in bacteria, yeast and mammalian cells ^8,10^. For example, yeast cells selected under low glucose conditions amplify the hexose transporters ^11^, and mammalian cells selected in methotrexate amplify the DHFR gene ^12–14^. Developmentally and environmentally induced gene amplifications gave rise to a model of adaptive amplification ^15^ that gene amplification events facilitated genetic adaptability, allowing cells to respond to changing environmental conditions. Initial experiments in bacteria provided evidence for the existence of directed gene amplification in response to environmental stress; however, whether gene amplifications in response to environmental cues are a directed cellular response or the outcome of stochastic events that are selected has remained elusive ^15^.

Previous work described transient site-specific gene amplification events (TSSGs) as an epigenetically regulated cellular program to induce DNA rereplication of specific genomic regions^17,19–22^. Human cells use TSSGs to respond to environmental cues (hypoxia) and growth factor signaling (EGF) by targeting epigenetic changes to specific genes resulting in a more open chromatin environment ^17,22^. The increased accessibility produces a targeted DNA rereplication event that generates an extrachromosomal piece of DNA. TSSGs are transient, as the extra gene copy is present only during S phase and early G2 and disappears prior to nuclear envelope breakdown ^19^. However, populations of cells retain the ability to reproduce TSSGs in subsequent cell cycles. TSSGs occur at regions frequently amplified in tumors including CKS1B and EGFR ^17,22^, suggesting that though initially transient, TSSGs may serve as precursors to stable, inherited gene amplifications ^17,19,21,23^.

We sought to determine if DNA rereplication events and TSSGs can act as precursors to inherited gene amplifications and provide a mechanism for adaptation to environmental stresses. We used a classical model of environmentally induced gene amplification, metal-induced amplification of the metallothionein genes (MT). MT genes encode small cysteine-rich proteins capable of binding to essential and non-essential metals ^24,25^. MT proteins are important for maintaining cellular metal homeostasis and have been identified in most organisms including prokaryotes.

Selected amplification of the MT genes in response to metals in Chinese Hamster Ovary (CHO) cells by unknown mechanisms was initially described by Hildebrand and colleagues in the 1980s ^26^. Long-term culture of CHO cells in increasing doses of cadmium selected a population with MT gene amplification. It has remained unclear how the selected MT amplifications originated and how specific the changes were to the MT locus. The importance of understanding MT amplifications has increased as MT genes are amplified and overexpressed in many tumors including breast, lung, brain, and bladder cancer ^24^ and higher MT expression correlates with poor prognosis and provides resistance to metal and non-metal-based chemotherapy ^27–30^.

To explore how cells respond to acute and chronic changes in environmental metal conditions, we developed cadmium-resistant breast cancer cells through continuous culture in increasing concentrations of cadmium. Cells initially respond through production of TSSGs of the MT locus that precede stable, inherited MT gene amplification. Cells normally suppress TSSG of the MT locus through trimethylation of histone 3 lysine 27 (H3K27me3) by the histone methyl transferase EZH2. Cadmium treatment results in loss of H3K27me3 near origins of DNA replication in the MT locus, which coincides with induction of DNA rereplication. Persistent ablation of EZH2 activity results in heritable MT gene amplification even in the absence of the selective pressure of cadmium. We demonstrate that inheritance of the MT gene amplifications requires the homologous recombination pathway (HR) and results in genomic structural variants (SVs) involving the MT locus and other chromosomal loci with homology to the MT locus. DNA rereplication induced in response to environmental metal exposure is conserved from yeast to human and adaptation to metal exposure requires origins of DNA replication in yeast. Adapted yeast cells also exhibit MT structural variants with chromosomal regions of microhomology to the MT locus. Our results describe a conserved DNA rereplication response to environmental stress that allows adaptation to changing environmental conditions through targeted, regulated gene amplification.

## RESULTS

### Human Cells Develop MT Amplifications in Response to Cadmium

MT gene amplification in response to long-term cadmium treatment has been well documented in rodent cells ^31,32^, however, it is unclear how MT amplification occurs in human cells and how the MT locus changes during development of metal resistance. Therefore, we adapted human cells to cadmium and monitored the development of MT-amplified, cadmium-resistant cells. We used breast cancer cells as a model because MT genes are frequently amplified in breast cancer patients (Figure S1a; ^27^), cadmium transforms normal breast cells ^33^, cadmium levels in breast tissue correlate with risk of cancer ^34^, and breast cancer cells have higher levels of cadmium compared to non-cancerous adjacent cells ^35^. MDA-MB-231 cells were specifically chosen as the MT amplification model cell line because they are diploid for chromosome 16 (Figure 1a).

**Figure 1:**
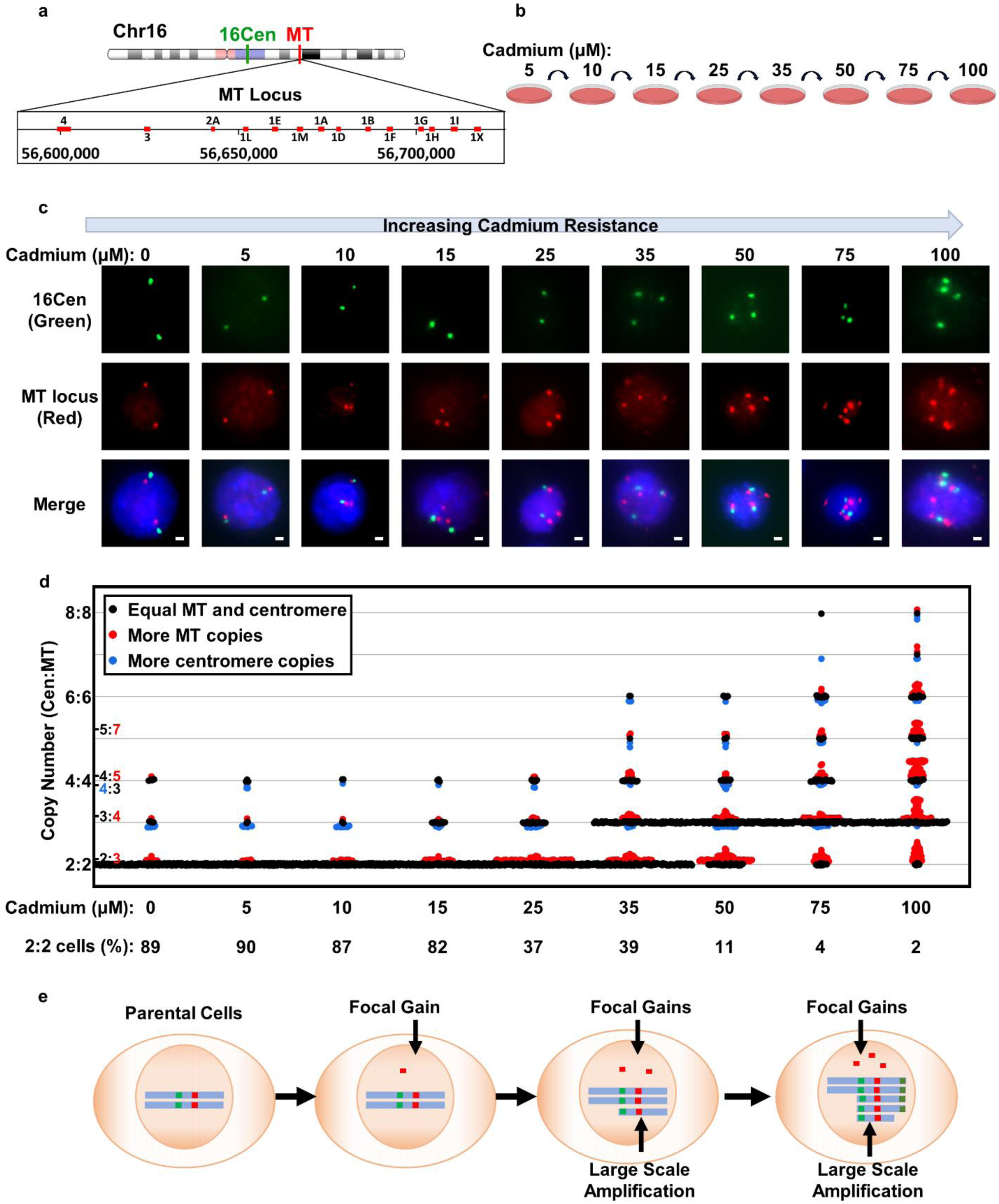
Cells Develop Progressive MT Amplifications in Response to Cadmium Exposure. (a) The metallothionein locus located on chromosome 16. (b) A schematic of cadmium treatment of MDA-MB-231 cells to develop cadmium resistant cell line. (c) Representative DNA-FISH images for cadmium treated cells at the indicated cadmium concentrations (d) Quantification of DNA FISH during development of cadmium resistance. Plot depicts 300 cells for each concentration (150 cells from each of 2 independent derivations of the resistant ceslls). Black dots represent cells with equal MT and 16Cen count. Red dots represent more MT foci than 16Cen. Blue dots represent more 16Cen than MT. (e) Schematic indicating change in MT copy number in cells. Scale bars represent 2 µM. See also Figure S1

To develop cadmium-resistant MDA-MB-231 cells (MDA-MB-231CadR), we treated cells chronically with cadmium (Figure 1b), starting at 5 μM and increasing dose every 10 passages as cells became tolerant to the increased cadmium concentration (see methods). We monitored the impact of cadmium on the MT locus using a custom-developed dual DNA-FISH probe. Our dual FISH probe contains a control probe (in green; 16Cen) that recognizes the alpha satellite centromeric heterochromatin of chromosome 16 approximately 15 megabases (Mb) upstream from the probe that recognizes the 150 kb MT locus (red; Figure 1a). Untreated, parental MDA-MB-231 cells have two foci for each probe (Figure 1c left), as do cells following treatment with low doses of cadmium (5 µM). However, as cells develop resistance to 10 µM cadmium, a fraction of the cell population exhibits three copies of the MT locus and two copies of the centromere indicating focal amplification of the MT locus (Figures 1d, 1e, S1d and S1e). Continued culture of cells in increasing concentrations of cadmium results in further increases in the copy number of the MT locus, and eventually an increase in the copy number of the centromeric alpha-satellite control probe (Figures 1d and 1e). When cells were able to tolerate 75 µM cadmium (MDA-MB-231CadR), greater than 90% of cells had amplified the MT locus. The MDA-MB-231CadR cells were a mix of larger-scale amplifications (increased red and green foci) and an increase in focal amplifications of the MT locus (more red foci than green foci). The population of MDA-MB-231CadR cells were non-clonal as many cells exhibited vastly different copy numbers of the MT and 16Cen foci (measured in the same cell at the same time), demonstrating that we did not isolate a small cohort of resistant clones.

The MDA-MB-231CadR cells are 40-fold more resistant to cadmium than the matched parental cells grown in parallel (Figure S1b), but also exhibit increased resistance to cisplatin (Figure S1c), to which MDA-MB-231 cells are normally highly sensitive ^36,37^. This cross-resistance is consistent with literature reports that increased MT expression provides resistance to cisplatin treatment in breast cancer cells ^38^. MT gene expression was increased with acute cadmium treatment (72 hours, Figure S1f) and continued to increase as cells developed cadmium resistance (Figure S1g). The timing of transcriptional changes was independent of the development of MT amplification events suggesting the potential for two separate mechanisms for adaptation.

### Cadmium Induces TSSG of the MT Locus

Our prediction was that cells would develop TSSG of the MT locus during the development of cadmium resistance. To test this hypothesis, we asked if at any stage during the development of MDA-MB-231CadR cells, the MT amplifications observed were transient. TSSGs are generated through DNA rereplication during S phase and thus disappear if the cell cycle is arrested ^19^. We arrested parental and cadmium treated cells in G1/S with hydroxyurea (HU; Figures 2a and S2A) and expected cells containing TSSG of the MT locus to exhibit a reduced number of cells with extra MT (red) foci. Indeed, HU treatment of cells in 10 μM cadmium blocked cadmium-induced MT focal (Figures 2b, 2c, 2e and S2a). Furthermore, treatment of MDA-MB-231CadR cells with HU resulted in a reduction of cells with extra MT foci (Figures 2d,2f and S2b). DNA-FISH of MDA-MB-231CadR cells using the MT locus probe with either the chr16 centromere probe or a chromosome 16 q-arm telomere probe (16-qtel) shows that the number of cells with MT and 16-qtel gains is much lower than cells from the same population with MT and 16Cen gains indicating that the amplifications do not represent whole chromosome nor arm level amplifications (Figures S2c and S2d). These results are consistent with the hypothesis that at low cadmium concentrations, cells respond with TSSG of the MT locus that eventually proceeds to stable, inherited MT gene amplifications. Furthermore, cells with stable inherited MT amplifications continue to produce some focal MT TSSGs. TSSG events in response to cadmium were specific to the MT locus as other regions known to undergo TSSG were not affected by cadmium treatment (Figure S2e).

**Figure 2:**
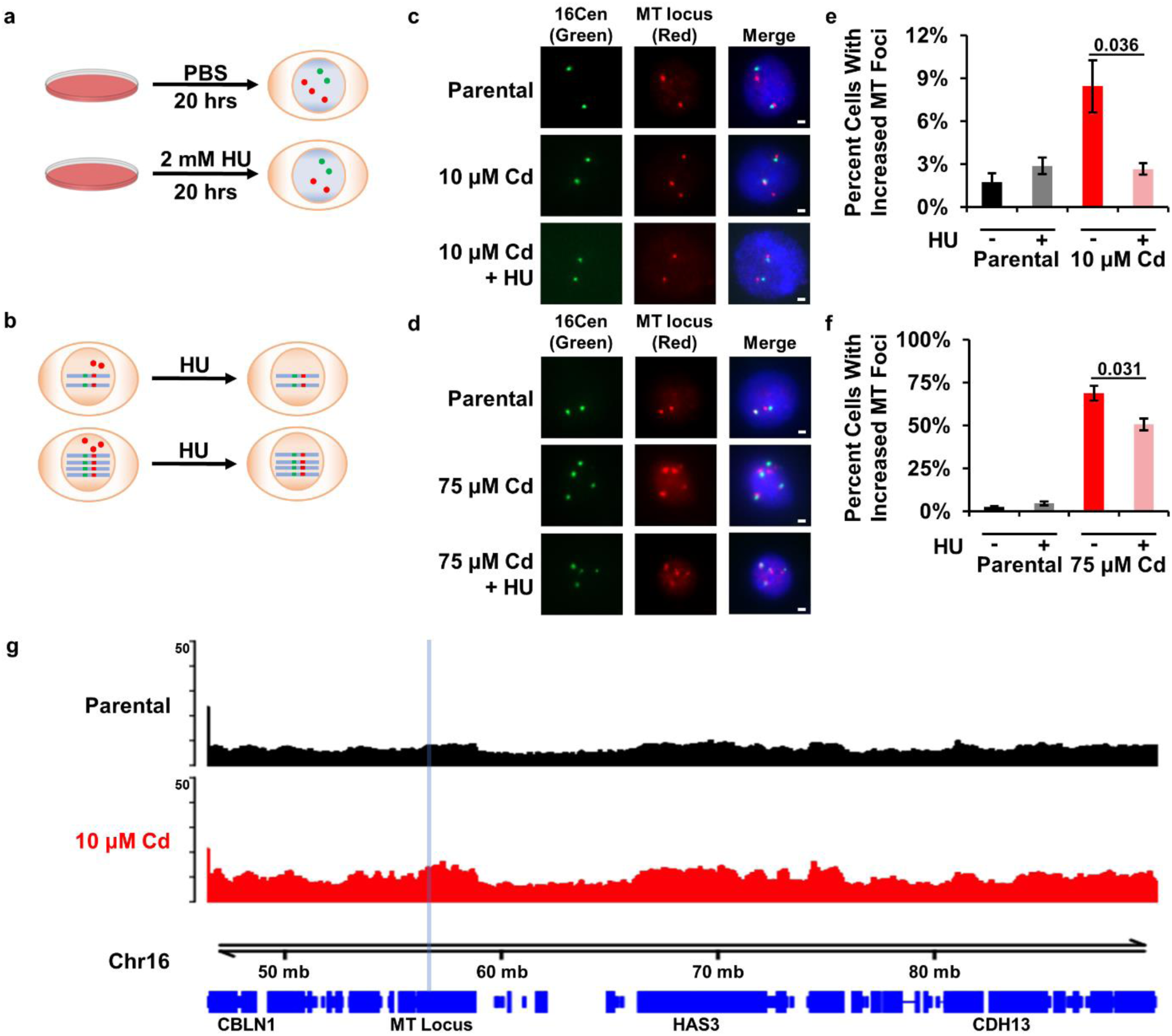
Cadmium Induces Transient Site-Specific Gains of the MT Locus. (a,b) Schematic of hydroxy urea (HU) treatment in MDA-MB-231 cells. (c,d) Representative DNA-FISH images for treatment of 10 µM and 75 µM Cd cells with HU. (e) 10 µM cadmium treated cells exhibit transient focal gains which are lost when treated with HU. (n=3 independent cell cultures). (f) 75 µM cadmium treated cells contain both focal and stable amplifications, only transient gains are lost when treated with HU. (n=3 independent cell cultures). (g) Rerep-seq analysis of 10 µM Cd treated cells shows that rereplication occurs throughout the MT locus and extends towards the chromosome 16 centromere. Rerep-seq tracks represent the mean of two biological replicates. Error bars represent the S.E.M. P-values determined using two-tailed Student’s t-test. Scale bars represent 2 µM. See also Figure S2.

### Cadmium Induces DNA Rereplication of the MT Locus

TSSGs are derived from DNA rereplication events ^17,19,21^. To determine if the MT locus undergoes DNA rereplication in MDA-MB-231CadR cells we used Rerep-seq to sequence rereplicated DNA ^39^. We performed Rerep-seq for both parental and MDA-MB-231CadR cells. Rerep-seq enzymatically cleaves BrdU incorporated during DNA replication to produce single strand breaks in normal replicated DNA, but staggered double strand breaks in rereplicated DNA, allowing it to shear and be isolated by size selection for analysis by qPCR or sequencing ^39^. We observed a significant increase in DNA rereplication throughout and surrounding the MT locus on chromosome 16 in both 10 µM and 75 µM cadmium treated cells (Figures 2g and S2f). This rereplication was specific to the MT locus as we did not identify other broad domains of rereplication at other genomic loci (data not shown).

Together our data indicate that cadmium treatment induces DNA rereplication and TSSG of the MT locus. Our results suggest that MT TSSGs may be precursors for inherited gene amplifications.

### EZH2 and H3K27me3 Suppress TSSG of the MT Locus

Previous examples of TSSGs were regulated by epigenetic changes that resulted in more open chromatin environments ^19,21,23,40^. The MT genes are prototypical EZH2 target genes. Depletion or genetic ablation of EZH2 (methyltransferase) or other components of PRC2 complex EED or SUZ12 results in transcriptional activation of the MT genes in mouse and human cells ^41–46^. Additionally, the MT locus is tri-methylated at H3K27 in mouse and human cells ^41–46^. Therefore, we reasoned the H3K27me3 and EZH2 activity may also regulate the development of MT TSSGs.

To address this hypothesis, we inhibited EZH2 methyltransferase activity using the inhibitor UNC1999 ^47^. Treatment of MDA-MB-231 cells with 810 nM UNC1999 for 72 hours resulted in a marked reduction of global H3K27me3 (Figure S3a) and an increased expression of the MT genes (Figure S3b). This short-term inhibition with UNC1999 resulted in an increase in focal MT copy number in parental MDA-MB-231 cells (Figures 3a, 3b and S3c) in as little as 24 hours. In further support of the transient nature of these amplifications, removal of UNC1999 for 24 hours was sufficient to ablate the MT amplifications (Figure S3c). In addition, the focal increase in MT copy number in MDA-MB-231 cells was suppressed by arresting cells with HU, indicating that inhibition of H3K27me3 results in TSSG of the MT locus (Figures 3a and 3c).

**Figure 3:**
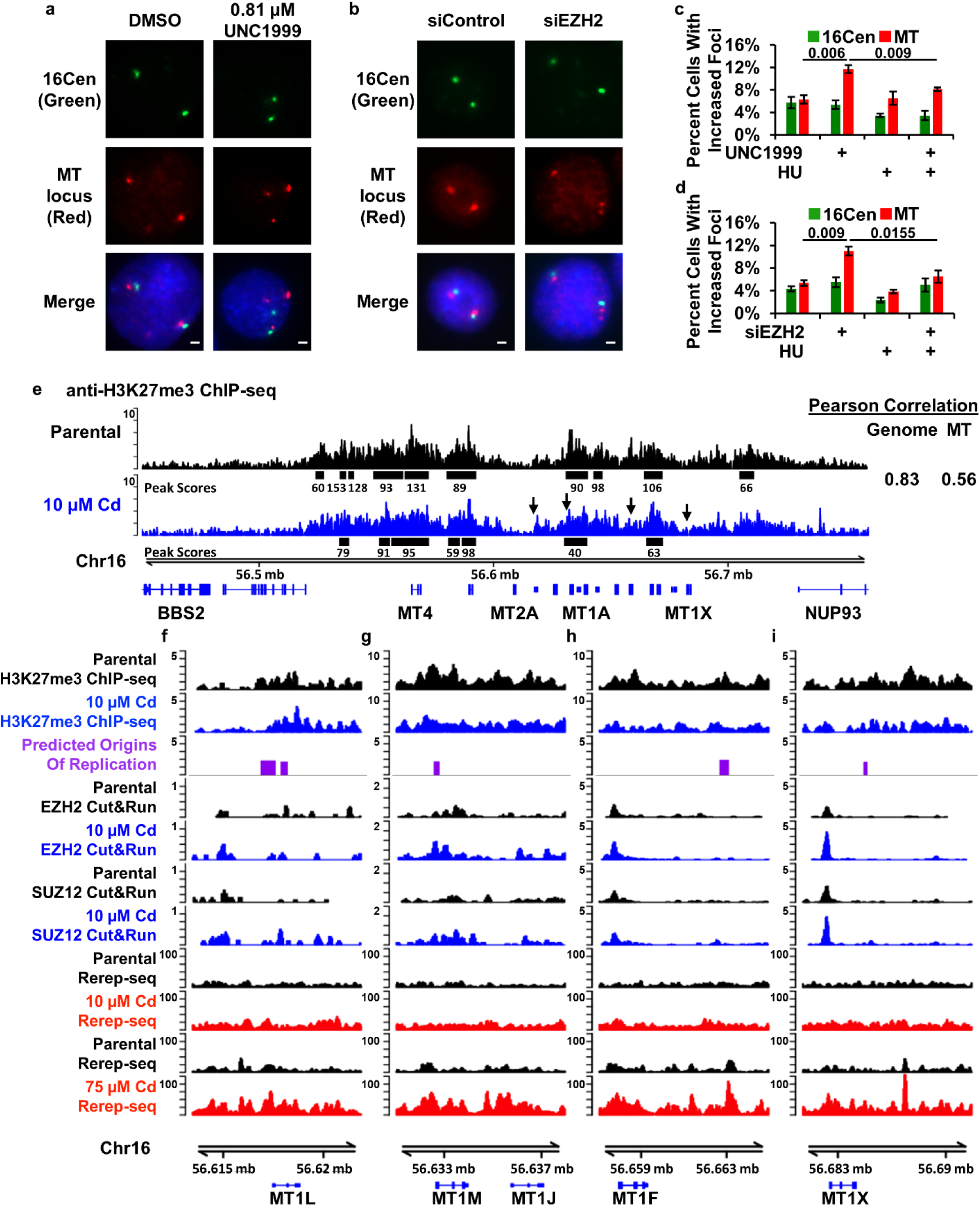
EZH2 and H3K27me3 Suppress TSSG of the MT Locus. (a) Representative DNA-FISH images for treatment of MDA-MB-231 cells with 810 nm of the EZH2 inhibitor UNC1999. (b) Representative DNA-FISH images for MDA-MB-231 following depletion of EZH2. (c,d) Inhibiting (c, n=3 independent cell cultures) or depleting (d, n=4, two independent transfections of two different siRNA) EZH2 promotes TSSG at the MT locus. (e) ChIP-seq demonstrates H3K27me3 is reduced at the MT locus in 10 µM Cd MDA-MB-231 cells. Black bars indicate peaks determined by Epic2 broad peak calling in H3K27me3 compared to input. Peak scores are indicated below called peaks and are decreased at peaks throughout the MT locus in 10 µM cadmium treated cells. Bin level Pearson correlation (right) indicates the H3K27me3 pattern correlates less well in the MT locus than throughout the genome. Black arrows indicate regions analyzed in f-i at higher resolution. (f-i) Loss of H3K27me3 signal correlates with increased rereplicated DNA from Rerep-seq. EZH2 and SUZ12 recruitment is equivalent or increased in cadmium treated cells. H3K27me3 ChIP-seq tracks represent the mean of 3 biological replicates, EZH2 and SUZ12 CUT&RUN and Rerep-seq tracks represent the mean of 2 biological replicates. Error bars represent the S.E.M. P-values determined using two-tailed Student’s t-test. Scale bars represent 2 µM. See also Figure S3.

To verify that induction of MT TSSG was an on-target effect of UNC1999, we depleted EZH2, SUZ12 and EED using siRNA (Figures S3d and S3e). Depletion of EZH2 resulted in increased focal amplification of the MT locus, which was suppressed by treatment with HU, indicating that loss of EZH2 induces TSSG of the MT locus (Figures 3b and 3d). In agreement with this, depletion of SUZ12 or EED also resulted in focal amplification of the MT locus (Figure S3h). These results were not specific to breast cancer cells, as depleting EZH2, SUZ12 and EED in immortalized, but non-transformed retinal pigment epithelial (RPE) cells also resulted in TSSG of the MT locus (Figures S3f, S3g, S3i and S3j). Taken together, our results demonstrate that H3K27me3 mediated by EZH2 regulates TSSG of the MT locus.

We investigated the levels of H3K27me3 at the MT locus by chromatin immunoprecipitation sequencing (ChIP-seq). Sequencing of these samples revealed a modest decrease in H3K27me3 throughout the entire MT locus (Figure 3e). Further, broad peak calling of H3K27me3 in the 10 µM Cd cells revealed a reduction in peak scores (reduced methylation within the peaks) in the cadmium treated cells compared to parental cells (Figure 3e black bars). Cadmium treatment did not result in a genome wide loss of H3K27me3 as the distribution of peak scores did not change significantly in cadmium treated cells relative to control cells (Figure S3k). We also did not observe a decrease in H3K27me3 at other highly H3K27me3 enriched loci, such as the CCND2 locus (Figure S3l).

To understand how changes in H3K27me3 might be related to DNA rereplication of the MT locus, we examined H3K27me3 ChIP-seq levels near known origins of replication in the MT locus in breast cancer cells ^48^. We observed loss of H3K27me3 at several locations in the MT locus that were at or adjacent to an origin of DNA replication (Figures 3F-3I). We then compared these locations in our Rerep-seq data and observed an increase in DNA rereplication surrounding regions that lost H3K27me3 signal. To determine if the reduction of H3K27me3 was caused by loss of EZH2 recruitment to these loci, we performed CUT&RUN ^49^ for EZH2 and SUZ12. Surprisingly, we observed increased EZH2 and SUZ12 occupancy at the MT locus, but not at CCND2 (Figures 3F-3I and S3L). This suggests that EZH2 activity in the MT locus is inhibited, or H3K27me3 is restricted near origins of DNA replication in the MT locus in cadmium treated cells. This is consistent with models of blocking methylation by EZH2 while it remains bound to target loci ^50,51^. Altogether, our results suggest that H3K27me3 mediated by EZH2 suppresses DNA rereplication in the MT locus and the loss of H3K27me3 correlates with regions of DNA rereplication.

### Prolonged EZH2 inhibition results in stable MT amplification without the selective pressure of cadmium

The MDA-MB-231CadR cells developed TSSG of the MT locus at low levels of cadmium and progressed to high level inherited gene amplification at higher concentrations of cadmium, suggesting that TSSG of the MT locus could precede the stable inherited gene amplification events. Notably, this was in the presence of cadmium, which provided increased selective pressure for maintaining the increased copy number of the MT genes. To determine if selective pressure was necessary for the transition from transient to inherited MT amplification, we examined the MT locus in cells chronically treated with the EZH2 inhibitor UNC1999. Initially, UNC1999 treated cells produced only focal transient MT gains (Figures 4a and 4b). However, after 20 to 30 passages in UNC1999, the cells began exhibiting increased copy number of the centromere and the MT locus (Figure 4a and 4b), characteristic of the inherited gains observed in cells cultured in cadmium (Figure 4c). The increased copy number in long term culture resulted in integrated MT copies observable in metaphase spreads suggesting these cells did indeed obtain inherited gene amplifications (Figure S4a). Long term culture of the cells in 10 µM cadmium and UNC1999, led to more rapid development of inherited MT gene amplification (passage 15-20; Figure 4d). Transcription of the MT genes was not greatly affected with UNC1999 treatment (Figure S4b), but was activated with the addition of cadmium (Figure S4c) consistent with the requirement for metal exposure to activate the MTF1 transcription factor ^52^. However, ablation of MTF1 in UNC1999 treated cells did abrogate production of MT TSSGs suggesting the DNA binding transcription factor may play a role in targeting the NDA rereplication response (Figure S4d and S4e). Combined, our results demonstrate that the selective pressure of cadmium is not required for stable inherited amplification of the MT locus, which suggests that continued generation of the MT TSSGs through environmental exposure to metal or deregulation of the epigenetic landscape, eventually allows cells to integrate and inherit the copy number alteration.

**Figure 4:**
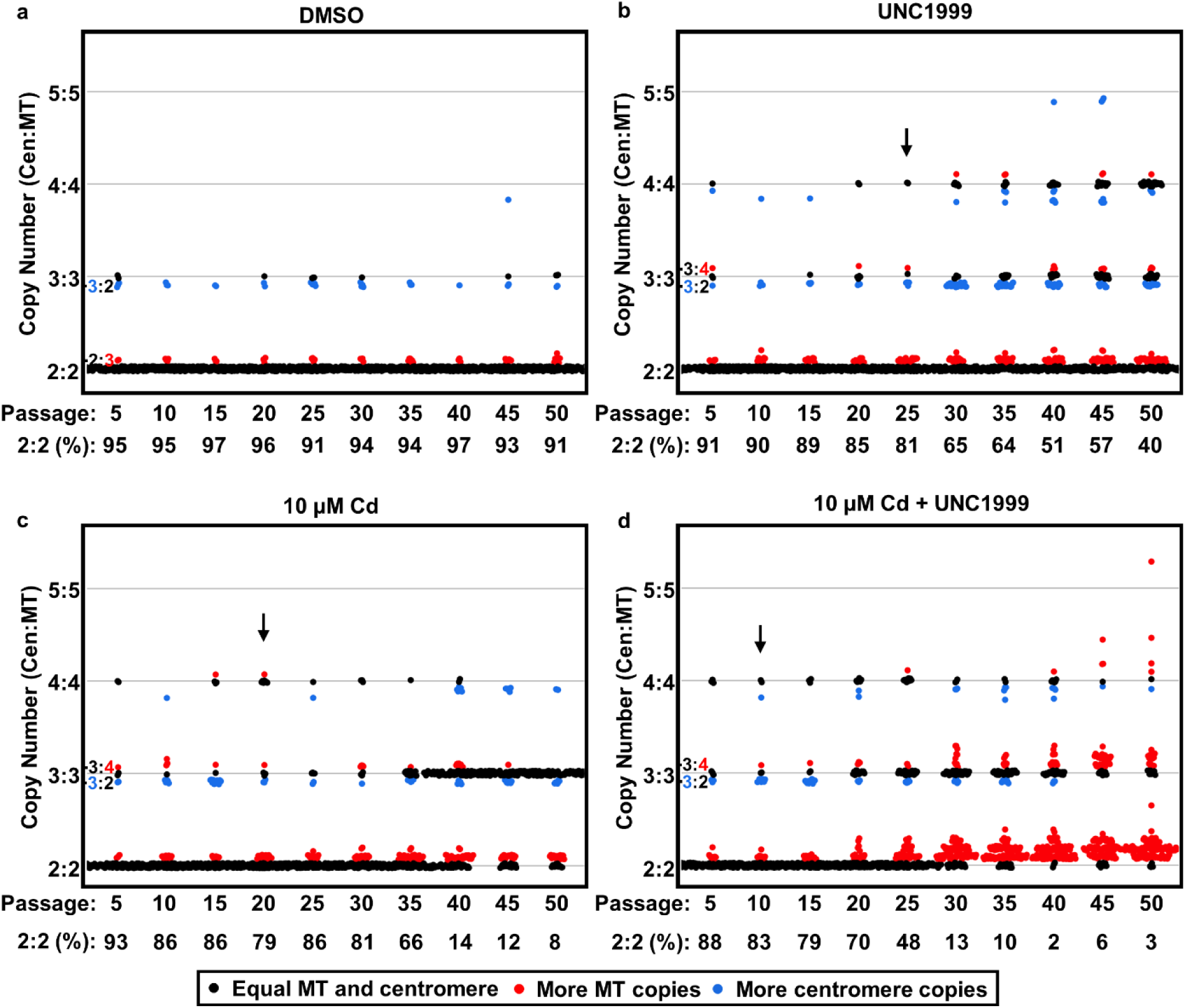
Prolonged EZH2 Inhibition Results in Inherited Metallothionein Amplification Without the Selective Pressure of Cadmium. (a-d) DNA FISH analysis for MT copy number following prolonged EZH2 inhibition. FISH analysis for MDA-MB-231 cells treated chronically with DMSO (a),810 nM UNC1999 (b), 10 µM Cd (c) or dual treatment with 810 nM UNC1999 and 10 µM Cd (d). Data represents 150 cells counted from each passage for each treatment condition. Black dots represent cells with equal MT and 16Cen count, Red dots more MT foci than 16Cen and Blue dots more 16Cen than MT. See also Figure S4.

### Homologous Recombination is Required for the Transition to Inherited MT Gene Amplification

We next determined how cells transition from TSSG to stable inherited MT gene amplification. We performed metaphase spreads to ascertain the state of MT amplifications in MDA-MB-231CadR cells. The majority of cells with stably integrated MT amplification exhibited repeat amplifications of the MT locus (Figure 5a), with some examples of integration on other chromosomes (MT foci not containing a 16Cen focus). Most current models of gene amplification involve homologous recombination ^53,54^ suggesting that HR is required for inheritance of the MT amplification. To test this hypothesis, we used long-term culture of cells in the presence of the EZH2 inhibitor UNC1999. We reasoned that performing this in the cadmium-derived MT amplified cells while blocking inheritance might result in sensitization of the cells to cadmium and the selective pressure of cadmium may bias the outcome. To determine if HR is required for inherited MT amplifications, we combined inhibition of EZH2 using UNC1999 with chemical inhibition of ATR using VE-821, which ablates the HR pathway within 6 passages (Figure S5a; ^55^). We performed this analysis using two different approaches: first, we added the HR inhibitor after 20 passages in UNC, right before the cells began integrating and inheriting the MT amplifications; and second, beginning a new culture with the HR inhibitor added to UNC from the beginning of the culture. In both experimental setups, inhibiting HR ablated the rise of stable, inherited MT amplifications (Figures 5b-5d and S5b-S5e).

**Figure 5:**
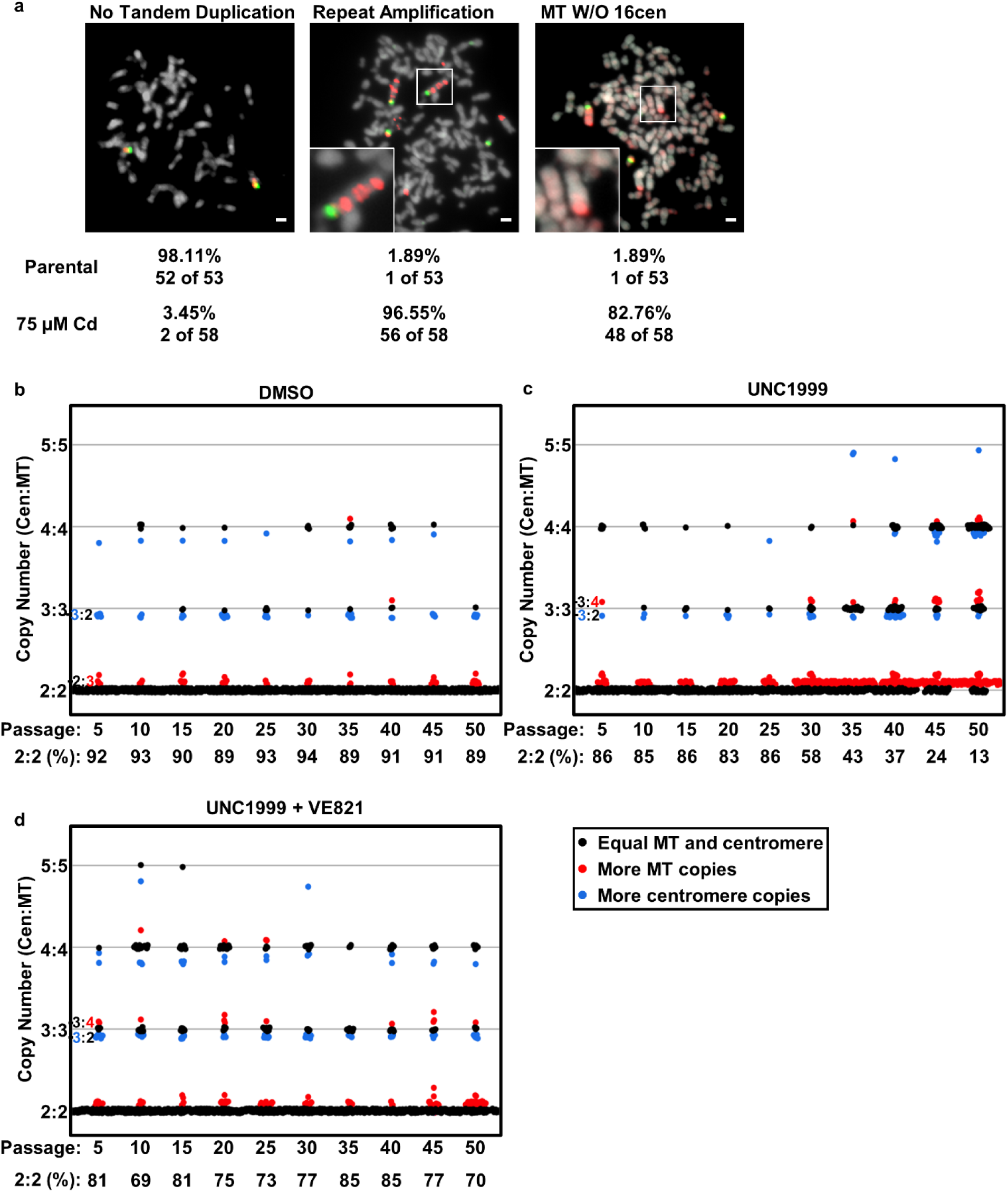
Homologous Recombination is Required For Inherited MT Amplification. (a) Representative DNA-FISH on metaphase spreads for parental and MDA-MB-231CadR cells. MDA-MB-231CadR cells display increased populations with repeat amplification and MT gains without chromosome 16 centromere. Numbers indicate percentage of cells scored that exhibited amplification of the indicated types (n>50 as indicated). (b-d) Inhibition of HR blocks inherited MT amplifications. FISH counts for cells treated with 810 nM of the EZH2 inhibitor UNC1999, of 5 µM of the homologous recombination inhibitor VE-821 or combined treatment. DNA-FISH data represents 150 cells counted from each passage for each treatment condition. Black dots represent cells with equal MT and 16Cen count. Red dots represent more MT foci than 16Cen. Blue dots represent more 16Cen than MT. Scale bars represent 2 µM. See also Figure S5.

To confirm these results in the cadmium adapted cells, we performed whole genome paired-end sequencing on the MDA-MB-231CadR cells and used GRIDDS to detect structural variants ^56^. We identified 191 variants within the MT locus consistent with the tandem duplications observed in our metaphase spread analyses (Figure 5a). Our metaphase spread analyses also suggested that MT amplifications could integrate on other chromosomes, so we used GRIDDS to identify SVs connecting the MT locus and a locus on another chromosome. We identified 485 variants that originated in the MT locus and connected to other chromosomal locations (Figure 6a). To determine if these regions were homologous to the MT locus, we identified the break point in the MT locus and the breakpoint at the other chromosomal locus and then retrieved the sequence within 500-bp up and downstream from the GRIDDS predicted break site. We used BLASTN (see methods) to align these two 1 kb segments and asked if there was homology between these regions. 370 of the 485 regions exhibited homology between the MT locus breakpoint and the other chromosomal region. This was significantly more than using random genomic regions or scrambling the same 1 kb sequences (Figures 6b-6e). These 370 regions with homology also had a lower E-value (better homology) and a longer average length of homology than the random genomic regions or scrambled sequences (Figure 6d and 6e), indicating the homology was due to the specific sequences and not just the nucleotide composition of the MT and insertion loci. We then averaged the homology at each position within the 1 kb fragment for all 485 alignments around the breakpoint (position 0) and observed significant enrichment of homology in the matched samples just upstream of the GRIDDS identified break point. We then analyzed the homologous sequences from each structural variant and identified that more than half contained repeat elements defined in the DFAM database ^57^. Compared to the genome-wide distribution of elements, we observed significant enrichment of several families of ALU repeat elements (Figure S6a), which is consistent with the idea that ALU elements can facilitate HR in human cells ^58,59^.

**Figure 6:**
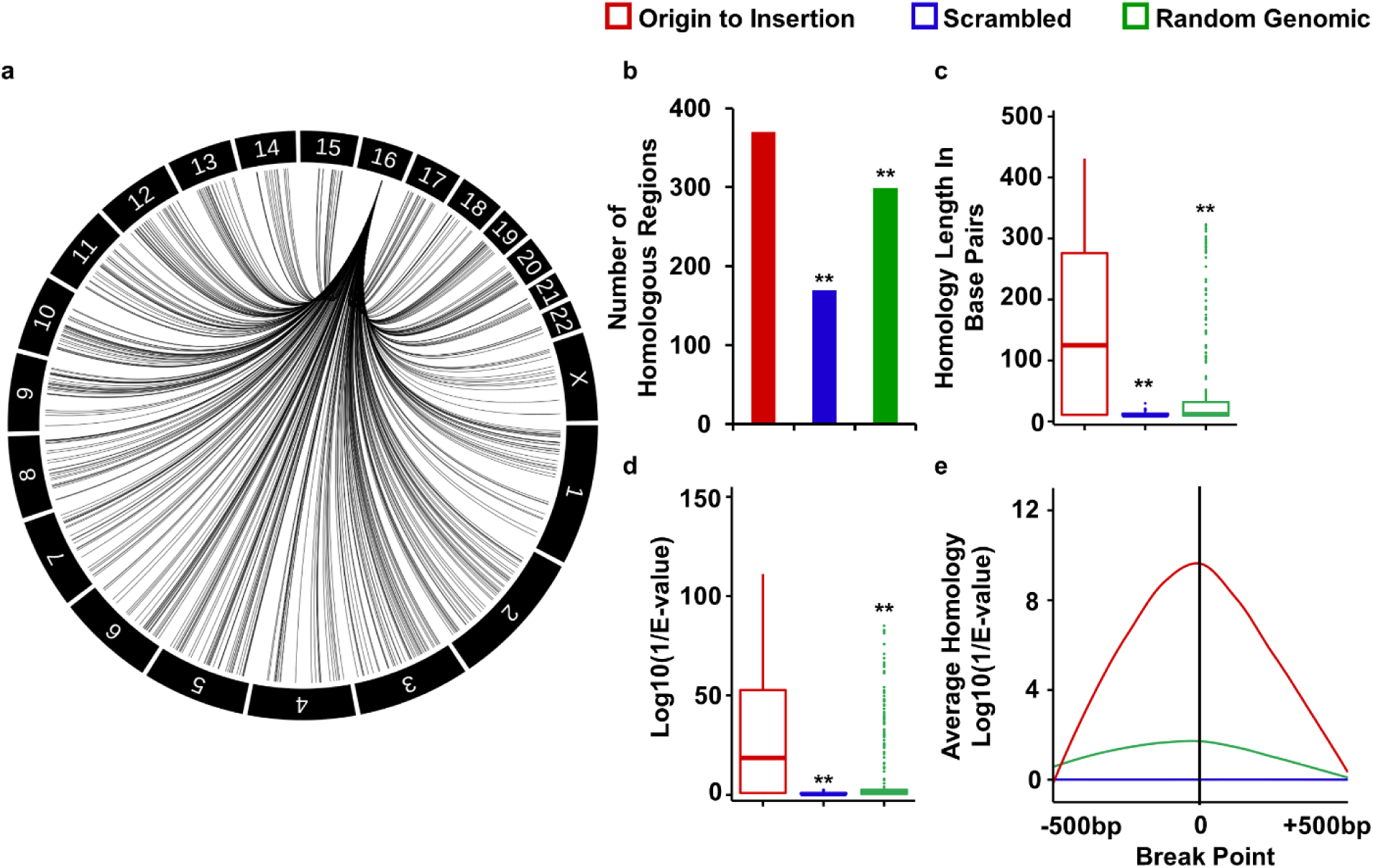
Identification of Interchromosomal Structural Variants with Homology to the MT locus. (a) Interchromosomal structural variants identified by GRIDDS analysis of MDA-MB-231CadR whole genome sequencing connecting the MT locus to locations throughout the genome. (b) Number of Interchromosomal structural variants with homology between the sequence surrounding the break point in the MT locus and the sequence surrounding the break point in the destination locus (Insertion). (c) MT-Insertion pairs have longer homologous regions compared to random genomic loci or the scrambled insertion sequence. (d) MT-Insertion pairs have better E-values compared to random genomic loci or the scrambled insertion sequence. (e) The homology in MT-insertion pairs is enriched immediately upstream of the GRIDDS predicted break points. ** indicates p < 1E-12 using a Kruskal–Wallis test with a post hoc Wilcoxon–Mann–Whitney test. See also Figure S6.

We analyzed the remaining 115 structural variants that did not exhibit extended homology between break points for microhomology (Figure S6b-S6e). We determined that 108 of the 115 remaining regions exhibited microhomology between the structural variant loci and the MT locus (compared to 65 in scrambled and 75 by random genomic comparison). The matched breakpoints had a longer average length of homology and lower E-value (better homology) than random genomic regions or scrambled sequences (Figure S6c and S6d).

### Metal-induced DNA Rereplication is a Conserved Response to Metal Exposure in Yeast

Metallothionein genes are conserved in most eukaryotes ^60,61^. If DNA rereplication is an important biological response to environmental metal exposure we would expect the response to be conserved in other organisms. To test this hypothesis, we sought to determine if metal induced MT amplifications were conserved in the yeast *S. cerevisiae*. The yeast metallothionein gene CUP1 increases in expression upon treatment with copper ^62,63^ and selection in copper-containing medium results in cells with increased copy of CUP1 ^63^. We developed a copper-resistant yeast strain by culturing cells in the presence of 6 mM copper for approximately 30 generations. Measurement of CUP1 genomic DNA (located on chromosome VIII) by qPCR demonstrated a significant site-specific increase in CUP1 DNA relative to the neighboring genes CIC1 and RSC30, and sub-telomeric locations on chromosome VIII (Figures 7a and S7a). To determine if the copper-adapted yeast culture was rereplicating CUP1, we performed Rerep-seq digests on the parental and copper-adapted cultures. We found that the CUP1 locus exhibited DNA rereplication in the copper-adapted culture (Figure S7b). We performed Rerep-seq on these cultures to determine the extent of the DNA rereplication of the CUP1 locus in the copper adapted strain. We observed strong rereplication of the CUP1 locus from 212,300 to 216,250 on chromosome VIII (Figure 7b). This is consistent with the boundaries of CUP1 amplicons from other studies ^64,65^. Our results indicate that yeast also undergo DNA rereplication of their MT locus in response to environmental metal exposure.

**Figure 7:**
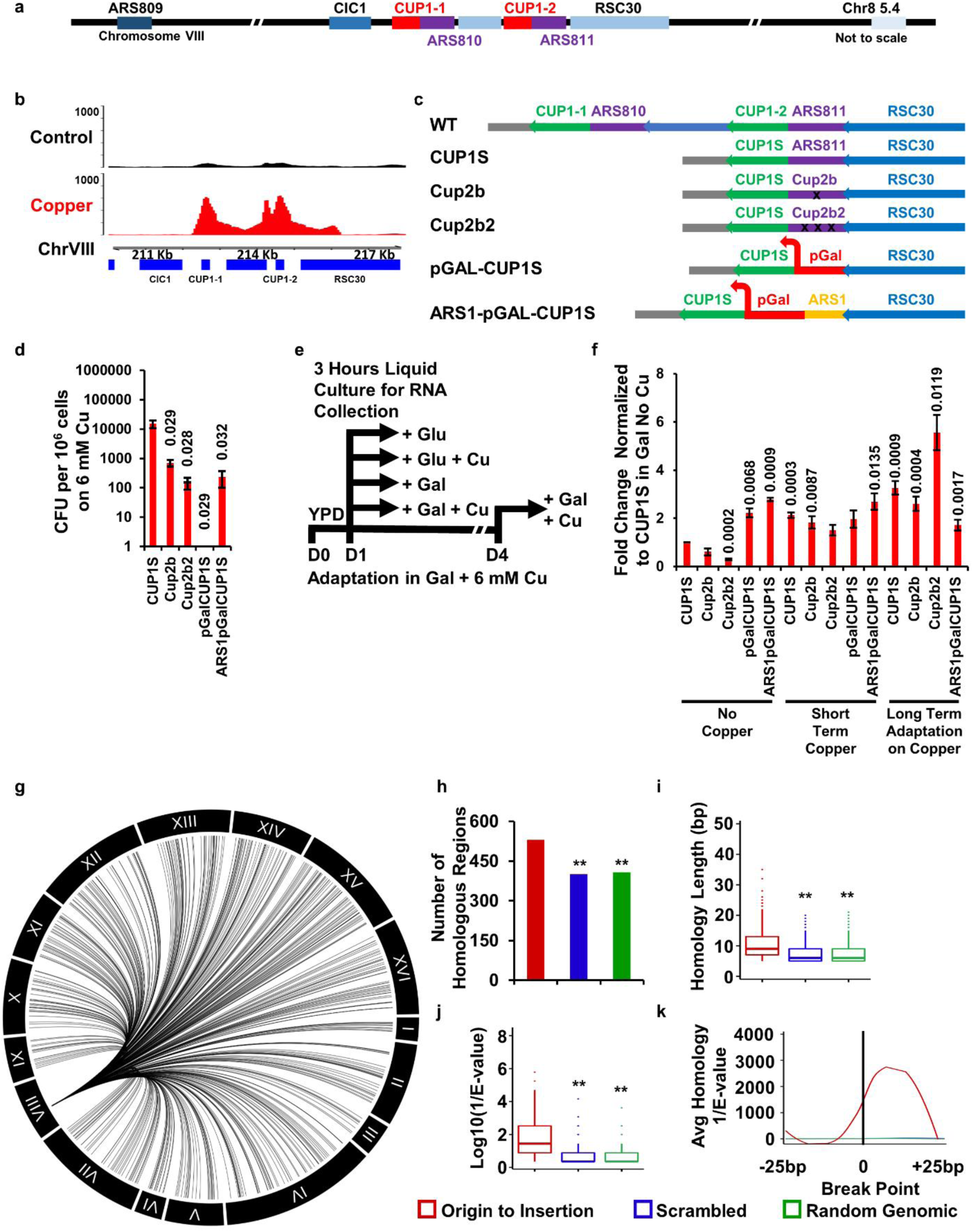
Metal-induced Metallothionein DNA Rereplication is Conserved in Yeast. (a) Schematic of the yeast chromosome 8 indicating locations of yeast metallothionein locus, containing CUP1 (in red). (b) Rerep-seq analysis of yeast grown in copper shows strong enrichment of the CUP1 locus. (c) Schematic of the yeast metallothionein locus and the strains engineered in this study. (d) An ARS is required for adaptation to copper. CFU of clones able to survive on YP 2% galactose plates supplemented with 6 mM copper. (n=3 independent clones). (e) Experimental approach to determine CUP1 mRNA expression from CUP1S mutants. (f) RT-qPCR from cultures grown as described in E. qPCR signals were normalized to ACT1 and related to the expression of Cup1S in 2% galactose without copper. Note pGAL-CUP1 does not adapt and does not grow on 6 mM copper and is hence absent from the selected/adapted set. (n=3 biological replicates). (g) Interchromosomal structural variants identified by GRIDDS analysis of copper resistant yeast whole genome sequencing. (h) Number of interchromosomal structural variants with microhomology between the sequence surrounding the break point in the MT locus and the sequence surrounding the break point in the destination locus (Insertion). (i) CUP1-Insertion pairs have longer microhomologous regions compared to random genomic loci or the scrambled insertion sequence. (j) CUP1-Insertion pairs have better E-values compared to random genomic loci or the scrambled insertion sequence. (k) The microhomology in CUP1-insertion pairs is enriched immediately downstream of the GRIDDS predicted break points. Error bars represent the S.E.M. (d,f) P-values determined using two-tailed Student’s t-test. (h-j) ** indicates p < 1E-12 using a Kruskal–Wallis test with a post hoc Wilcoxon–Mann–Whitney test. See also Figure S7.

### Molecular Determinants of Adaptation to Copper

The conservation of DNA rereplication as a response to environmental stress in yeast enabled use of yeast genetics to understand the molecular determinants of CUP1 amplification. Common laboratory strains of *S. Cerevisiae* contain two identical copies of CUP1, CUP1-1 and CUP1-2 with a partial duplication of RSC30/YHR056C (Figure 7c). In addition, this CUP1 locus is typically tandem duplicated, meaning most laboratory strains contain at least four copies of the CUP1 gene ^66^. The CUP1 promoter region contains consensus binding sequences for the yeast metallothionein transcription factor, Cup2. However, these elements are intermingled with an Autonomously Replicating Sequence (ARS811), a sequence-defined origin of DNA replication in *S. cerevisiae*. We therefore constructed several mutant strains to investigate the importance and contribution of the ARS and CUP2 binding elements to adaptation to high environmental copper.

To simplify interpretation of these mutations, we first constructed a strain with a single copy of CUP1 by removing YHR054C (the partial RSC-30 duplication) and CUP1-1, including its promoter region and ARS810 (Figure 7c CUP1S,). This strain now has a single copy of CUP1 (the original CUP1-2). We then introduced a single point mutation (Figure 7c; Cup2b) or multiple point mutations (Figure 7c; Cup2b2) within the CUP2 binding motif ^67^; however, these mutations may also negatively impact the function of ARS811 as several ARS and CUP2 binding consensus sequences overlap. To circumvent this complication and distinguish between transcriptional and ARS function, we constructed two additional strains where we first replaced the entire CUP1 promoter region with the galactose inducible promoter (Figure 7c pGAL-CUP1S) and a second strain with a combination of the galactose inducible promoter with an upstream ARS1, a highly efficient early firing ARS from chromosome IV (Figure 7c ARS1-pGAL-CUP1S). Reducing the WT strain to a single copy of CUP1 resulted in approximately 10-fold less efficient adaptation to copper (Figures 7d-7f) on glucose containing media. Comparing the mutated strains to CUP1S, adaptation to copper was further diminished in Cup2b, and Cup2b2. The orthologous promoter strains, pGAL-CUP1S and ARS1-pGAL-CUP1S, failed to grow on the plates containing copper and glucose, which does not allow expression of CUP1 from the pGAL promoter (Figure S7c and S7d).

To better quantify adaptation to copper, we plated cells on a relatively high concentration of copper (6 mM) and 2% galactose, and counted surviving Colony Forming Units (CFU) (Figure 7d). Compared to CUP1S, Cup2b and Cup2b2 displayed diminished survival and produced approximately 10-fold and 100-fold fewer surviving colonies respectively (Figure 7d). pGAL-CUP1S was not able to adapt to copper and did not produce any colonies; while ARS1-pGAL-CUP1S was able to adapt to 6 mM copper, demonstrating the presence of the adjacent ARS1 was critical for adaptation (Figure 7d). We measured gene expression from CUP1 after short-term copper exposure (3 hours) to determine CUP1 mRNA levels at the beginning of the adaptation experiment, and from cultures from colonies at the end of the adaptation experiment (Figure 7e). Unexpectedly, Cup2b mutants did not significantly reduce CUP1 mRNA expression, and had increased expression after adaptation (Figure 7f), suggesting that mRNA production does not account for reduced survival, and that these mutations perhaps diminished the function of the overlapping ARS811. Prior to adaptation, pGAL-CUP1S and ARS-pGAL-CUP1S express levels of CUP1 similar to CUP1S in galactose and copper. However, post adaptation, ARS1-pGAL-CUP1 expressed lower levels of CUP1 than CUP1S.These data indicate that transcription of the CUP1 gene was not sufficient to allow adaptation to copper, but that adaptation requires the presence of an origin of replication, either a still functional ARS811 or the orthologous ARS1. Taken together, our results demonstrate that a DNA replication origin is required for CUP1S to adapt to high copper, which involves eventual reduplication of the CUP1 gene. Our results support a role for a replicative mechanism in metallothionein gene amplification and adaptation to high concentrations of copper.

Our experiments in human cells indicated that long-term adaptation to environmental metal led to inherited gene amplification. Consistent with this idea, genomic PCR using inverse primers extending away from one another, we found that copper adapted CUP1S yeast had reestablished a tandem repeat of the CUP1 gene (Figure S7e). To address how CUP1 amplification occurred in yeast, we analyzed structural variants from the CUP1 locus to other chromosomal locations. We performed GRIDDS analysis using paired-end whole genome sequencing in the copper adapted yeast strain. GRIDDS identified 10289 structural variants within the CUP1 locus and 558 variants linked to other chromosomal loci (Figure 7g). Using the same BLASTN procedure as in the human cells we did not identify significant long stretches of homology in the yeast data (data not shown). However, microhomology mediated recombination (MHMR) is frequently used in S. cerevisiae ^68,69^. Therefore, we adapted the BLAST procedure to identify microhomologies (4 bp or larger) within 50bp (25bp up or downstream) of the GRIDDS predicted break point. We identified 531 of the 558 regions with significant microhomology using this analysis (Figure 7h), which was greater than we observed using random genomic regions or scrambling the same sequences (407 or 401 sequences). The matched breakpoints had a longer average length of homology (10.68 vs 7.44 or 7.36) and lower E-value (better homology) than random genomic regions or scrambled sequences (Figures 7i and 7j). Finally, analysis of the position of the alignment showed a strong enrichment just downstream of the GRIDDs predicted breakpoint (Figure 7k), consistent with the position in MHMR models ^68,69^.

GRIDSS analysis of our Rerep-seq data revealed substantially fewer structural variants (Figures S7f and S7g; 55 in human and 80 in yeast). None of these variants overlapped with variants identified in the whole genome sequencing. This suggests that some variants may be rereplicated, but those variants integrated at other genomic locations are not being rereplicated at a detectable level.

Recent work has identified the widespread presence of extrachromosomal circular DNA (eccDNA) and indicated the importance of extrachromosomal circles in oncogene amplification ^70–73^. How eccDNA is generated is unclear, however, we reasoned that DNA rereplication or resolution of DNA rereplication could generate eccDNA. Consistent with this idea, we identified inverted read pairs in both our yeast and human whole genome sequencing analysis. These read pairs are consistent with the presence of eccDNA containing the MT and CUP1 locus. In human cells, we observed reads consistent with both large circles spanning 1-2 megabases (Figure S7h) and small circles coincident with the replication domain containing the MT locus (Figure S7i). In yeast, we observed a strong abundance of circles on the immediate boundaries of the CUP1 locus (Figures S7j and S7k) consistent with previous reports of highly defined circles from CIRC-seq data at the CUP1 locus ^64,74^.

The consistent finding of homologies across two species supports a conserved model whereby DNA rereplication in response to environmental metal transitions to stable, inherited gene amplification events through an HR mediated process.

## DISCUSSION

We have described an evolutionarily conserved DNA rereplication response to environmental stress that can transition to stable, inherited genetic changes. This new paradigm provides a mechanistic basis for how cells respond to acute stress that allows for eventual adaptation to chronic stress. Using the response to changes in environmental metal as an example, we demonstrate that following initial metal exposure, cells rereplicate their metallothionein genes. Continued exposure and adaptation to metal stress eventually results in stable MT gene amplification through an HR-dependent mechanism. Our results describe how transient site-specific gene amplification events can become stable, inherited gene amplifications. These findings have profound implications for understanding evolutionary gene duplication events, the function of stress response pathways, adaptation to acute and chronic stress, and the development of gene amplifications and tumor heterogeneity in cancer.

Previous work demonstrated that cells respond to hypoxia by inducing DNA rereplication of specific gene and non-gene targets ^17^. However, the targets that underwent DNA rereplication had no known relationship to survival or adaptation to the stress events. In addition, work from Clarke and colleagues demonstrated that EGFR amplifications exist in both a transient and permanent state and that increased EGFR protein level was required for augmented growth ^22^. In the present study, we demonstrate that DNA rereplication can amplify the MT genes in response to metal exposure, which the MT genes are required to adapt to ^61^. It is tempting to speculate that DNA rereplication and TSSG formation could be important under other stress response conditions where increasing gene dosage of key response genes provides a selective or growth advantage. It will be important in future work to determine how often DNA rereplication is used as part of stress responses and what signaling, and regulatory pathways dictate its use.

Adaptive amplification has been proposed as a mechanism for organisms or cells to cope with changing environmental conditions arose. This model from experiments in bacteria that demonstrated that amplification of the *lac* operon allowed rapid evolution to growth in lactose stress conditions ^75^. DNA amplifications and gene duplications have long been recognized as responses to environmental stress conditions and are often apparent in genes that provide selective pressure under stressful growth conditions ^76^. Adaptive amplification is favorable to adaptive mutagenesis as it does not involve mutation of the heritable gene copies that may or may not be favorable to growth in stress conditions. By amplifying specific genes, cells can respond to the environmental condition without damaging the original gene copy.

Here we provide evidence for adaptive amplification through DNA rereplication in human cells. Our results demonstrate that at least the initial transient response is directed and can be controlled by cells to respond to the environment. The use of DNA rereplication or TSSG in response to environmental stimuli provides an even greater advantage as the initial extra gene copies are transient and easily removed when the environmental stress disappears. However, prolonged stress exposure can provide conditions for either selection, through positive selective pressure, or stochastic integration at homologous sites. This would allow for eventual adaptation through heritable changes in gene copy number. Thus, DNA rereplication programs in response to stress could contribute to the fitness of an organism by allowing improved survival of somatic cells in response to environmental insults.

It has become apparent that extrachromosomal DNA is abundant in cell lines and in primary samples from cancer patients ^70–72,77–79^. Recent work has determined that at least some of this extrachromosomal DNA exists as circles and that these circles often contain amplified oncogenes ^74^. However, the origins of the extrachromosomal circular DNA (eccDNA) remain unknown. Several models propose that these circles can be excised from genomic DNA through homologous recombination or through circularization of broken chromosome fragments from mitotic defects ^80–82^. We propose that targeted DNA rereplication of the endogenous locus could be resolved, in part, through the production of circular DNA. In support of this idea, we identified reverse-forward inverted read pairs in our whole genome sequencing of yeast and human metal treated samples that are consistent with the presence of extrachromosomal circles (Figures S7h and S7i), though we note that these were rare events in the human cell models. This suggests that DNA rereplication could also be a source of circular extrachromosomal fragments. Consistent with this idea, the CUP1 locus in yeast is known to exist as eccDNA ^64^ and the boundaries identified in circular sequencing experiments from Møller and colleagues agree with the boundaries of our Rerep-seq data. It will be important in future studies to determine if DNA rereplication can be a source for eccDNA and how circular or linear extrachromosomal DNA production contributes to environmental stress responses.

Sites of DNA rereplication and TSSG regulated by the histone demethylase KDM4A occur at regions frequently amplified in tumors ^17,19,20,22^. Indeed, TCGA data can be used to predict TSSG sites for KDM4A by analyzing regions coamplified with KDM4A in tumors ^19^. This suggests that DNA rereplication and TSSG events could be precursors to gene amplifications observed in tumors. The epidermal growth factor receptor (EGFR) was recently reported to undergo TSSG formation ^23^. While it is not yet clear if the TSSG of EGFR can eventually become integrated and inherited, this implies that DNA rereplication events could be one origin for integrated and inherited oncogene amplifications. In addition, this suggests that some oncogene amplifications may not be the result of selection of random gene amplification events, but instead arise from targeted transient amplification of oncogenes. Through understanding the mechanisms regulating TSSG and the transition to inherited gene amplifications, it may be possible to prevent structural changes that could be induced by an EZH2 inhibitor or chemotherapy use in cancer patients and prevent structural changes that confer resistance to treatment.

The conserved nature of the MT amplification in S. cerevisiae CUP1 allowed us to perform analyses of the requirements for adaptation to copper in yeast. Surprisingly, this resulted in the observation that simply increasing transcription of CUP1 was not sufficient to allow yeast to adapt and survive in the presence of high levels of copper. Adaptation required the presence of an origin of DNA replication, though not necessarily the origin normally associated with CUP1. This data demonstrates that a replication event, and likely the rereplication of the CUP1 locus is required for adaptation to copper. It is important to note that removing the ARS from the locus did not cause growth defects on non-copper containing media indicating that this was a specific requirement in the presence of copper.

While transcription of CUP1 was not sufficient for adaptation, it remains unclear if transcription is required for DNA rereplication or TSSG generation. In the present study, the kinetics and magnitude of transcriptional changes are decoupled from TSSG formation. In cadmium exposed cells, transcription of the MT genes increased 5 to 10-fold within three days of metal exposure. However, the TSSGs do not develop until the cells are exposed to cadmium at to 10 µM roughly 7-14 days after beginning cadmium treatment at 5 µM, even though the transcriptional changes occurred within 72 hours of acute treatment (Figure S1f). In contrast, TSSGs rapidly develop when EZH2 is inhibited in 24 to 72 hours coincident with modest, acute transcriptional changes that do not persist in long-term EZH2 inhibition (Figures 3c, S3b and S4b). Thus, the appearance of the MT TSSGs is not correlated with the timing or magnitude of MT transcriptional induction. Transcription of the locus may still be important for initiation or resolution of DNA rereplication; however, more complex detailed experiments will need to be conducted to dissect out the requirement for transcription or the transcription factors, which may influence DNA replication.

In this manuscript, we describe targeted, regulated DNA rereplication as a conserved response to environmental stress. While initially transient, the rereplicated regions can become stably integrated and inherited through homologous recombination. Our results establish a new paradigm for the origin of gene duplication events, gene amplification and how cells can respond and adapt to environmental conditions.

## AUTHOR CONTRIBUTIONS

G.W., J.C.B and J.M. designed and conceived the study. G.W. and J.M performed the experiments. G.W. and P.T. performed bioinformatics analysis.

## ACKNOWLEDGEMENTS

This work was supported by NIGMS 1R35GM128720-01 to JCB. JCB is a Boettcher Foundation Scholar. We thank Jim Costello, James DeGregori, Patricia Ernst, Heide Ford, Paul Jedlicka, Andrew Thorburn, Capucine Van Rechem, Kelly Sullivan and Jonathan Whetstine

## DECLARATION OF INTERESTS

The authors declare no competing interests.

## EXPERIMENTAL PROCEDURES

### LEAD CONTACT FOR REAGENT AND RESOURCE SHARING

Requests for reagents and resources may be directed to the Lead Contact, Joshua Black

### EXPERIMENTAL MODEL AND SUBJECT DETAILS

#### Cell Culture

MDA-MB-231 female breast cancer cell line and hTERT-RPE-1 female retinal pigment epithelial cell line (called RPE throughout) were maintained at 37°C in Dulbecco’s Modified Essential Medium (DMEM) with 10% FB essence (VWR), 1% penicillin/streptomycin, and L-glutamine. Cadmium-resistant (MDA-MB-231CadR) cell lines were selected by serial passaging in medium containing CdSO_4_. The CdSO_4_ concentration was increased by increments of 5-25 μM after 10 passages at the current concentration and when cell count increased at least 1.8 times average daily over at least four passages consistently. Cells used in this study were adapted to CdSO_4_ containing medium for 7 months up to 100 μM CdSO_4_.

#### Yeast Strains, Plasmids, Media and Growth Conditions

Yeast strains and plasmids used in this study are listed in Table 1 and were grown according to standard procedures in YPD (yeast extract, peptone, and 2% dextrose) at 30°C for all experiments unless otherwise stated. To accommodate *S. cerevisiae*’s lack of the thymidine salvage pathway, yeast strains in this paper are all stably transformed with NheI linearized p403–BrdU–Inc HIS3, a reconstituted thymidine salvage pathway cassette, consisting of Herpes simplex virus thymidine kinase (HSV–TK) and human equilibrative nucleoside transporter (hENT1), that enables efficient cellular uptake and incorporation of the thymidine analogue BrdU into DNA ^83^. Inserts were confirmed by spot assays for sensitivity to 75 µg/ml FUDR and susceptibility to BrdU uptake and genomic fragmentation via Rerep-seq.

**Table 1.** *S. cerevisiae* strains

| Yeast Strain | Genotype | Background/Reference |
| --- | --- | --- |
| YJB1 WT | MATa ade2-1 his3-11,15 leu2-3,112 trp1-1 ura3-1 can1-100 bar1::hisG ars608Δ::HIS3 ars609Δ::TRP1 ars305::TRP1 GPD-HSV-TK ADH1-hENT1 BrdU-Inc | W303a/<sup>84</sup> |
| YJB21<br>*cup1Δrsc30Δ* | CUP1.1Δ C-terminal RSC30Δ::Ca URA3 | YJB1/(This study) |
| YJB26 CUP1S | *cup1Δrsc30Δ::CUP1-ARS811-RSC30* (single copy CUP1 with single ARS811 and restored C-terminal RSC30) | YJB21/(This study) |
| YJB30 *cup1Δ* | *cup1Δrsc30Δ::RSC30* (restored C-terminal RSC30 with No CUP1 or ARS810 or 811) | YJB21/(This study) |
| YJB31 Cup2b | ARS811::ARS811-CUP2 binding mutation | YJB26/(This study) |
| YJB36 Cup2b2 | ARS811::ARS811-CUP2 multiple binding mutations | YJB26/(This study) |
| YJB33 pGAL-CUP1S | CUP1S, ARS811Δ::pGAL | YJB26/(This study) |
| YJB41 ARS1-pGAL-CUP1S | ARS811Δ::ARS1-pGAL | YJB26/(This study) |

### METHOD DETAILS

#### Fluorescent In Situ Hybridization (FISH)

Cells were washed twice with PBS and treated with 80 mM potassium chloride for 15 minutes at 37°C. Cells were fixed in and washed twice with methanol-acetic acid (3:1). 100 μL of cells were added per well on an 8 well FISH slide and centrifuged 900 rpm for 3 minutes. The slide was incubated at room temperature (RT) in darkness for 10 minutes prior to washing. Slide was washed for 2 minutes in 2x SSC, 15 minutes in 0.01 M HCl with 50ppm pepsin, 5 minutes in PBS, 2 minutes each in 70% ethanol, 85% ethanol and 95% ethanol. Slides were dried for 10 minutes at room temperature. FISH was performed using probes for the MT locus in 16q12.2 (56,396,107 to 56,782,063 on chromosome 16 in GRCh37) and chromosome 16 α-satellite purchased from Oxford Gene Technology, who performed paired-end sequencing to verify the identity of the BACs. Probes were verified in RPE cells known to be diploid for both foci. Denaturation was performed at 78°C for 4 minutes and hybridization at 37°C 20-24 hours. Slide was washed in 0.4xSSC for 1.5 minutes at 72°C followed by 2XSSC with 0.05% tween 20 for 2 minutes at room temperature. Slide was incubated in 10 g/L BSA in 1X PBS with 0.5 µg/mL DAPI in PBS with 5% BSA for 5 minutes followed by 1X PBS for 5 minutes. Prolong anti-fade GOLD mounting medium (Molecular Probes, Life Technologies) was added to each well and sealed with a cover slide. Slide was cured overnight before imaging. Images of 20 planes in 0.5 µm steps for fields of nuclei were acquired on a Ziess Axio Imager M.2 microscope with an Orca-Flash 4.0LT digital camera and analyzed using Zeiss Zen2 software. We used a conservative scoring metric for copy gains. Any foci that were touching in the same focal plane were scored as a single focus to prevent increased numbers due to normally replicated foci. At least 100 cells for each of at least two biological replicates were scored for all experiments. FISH scores below 2 or above 9 were displayed as 2 or 9 respectively for the generated dot plots, though these were rare occurrences.

#### Metaphase Spread

MDA-MB-231CadR Cells were arrested in metaphase with 1 μg/mL colcimide treatment 1 hour prior to collecting as stated in FISH methods. Fixed cells in methanol-acetic acid (3:1) were dropped from 1-2 feet onto a glass slide above the water in a 55°C water bath at a 45-degree angle with the suspension allowed to run down the slide for 10 minutes with the slide maintained above the water in the water bath. Slide was then prepared as stated in FISH methods starting with 10-minute incubation before initial 2x SSC wash.

#### DNA Extraction

Parental and CadR cells were collected in PBS, resuspended in 1 mL modified RIPA buffer (50 mM Tris pH 7.4, 150 mM NaCl, 1% NP-40, 0.25% Sodium Deoxycholate, 1 mM EDTA pH 8.0, 20% glycerol). The samples were sonicated for 15 minutes using 30 second on/off cycle repeats. 2 μL RNaseA was added at 37°C for 2 hours. 10% SDS was added at 0.8% of the reaction volume as well as 10 μL Protease K (PK) and incubated at 55°C overnight. DNA was extracted with phenol, followed by addition of 1/9 reaction volumes of 3M sodium acetate and 70% reaction volumes of isopropanol. Samples were spun down at maximum speed for 17 minutes, washed with 750 μL 70% ethanol, spun down at maximum speed for 10 minutes, air dried 10 minutes and resuspended in 100 μL NF water. DNA concentration and purity were determined with a ThermoScientific Nanodrop 2000 Spectrophotometer measuring absorbance at 260, 280 and 230 nm.

#### Total RNA Extraction

Total RNA was extracted from parental and cadmium cells using the miRNeasy Mini Kit (Qiagen) following the manufacturer’s protocol including the optional DNase digest. RNA concentration and purity were determined with a ThermoScientific Nanodrop 2000 Spectrophotometer.

#### Acid Extraction of Histones

Cells were collected in PBS, washed in 5 mL PBS and resuspended in hypotonic lysis buffer (10 mM Tris-Cl pH 8.0, 1 mM MgCl2, 1 mM DTT). Resuspended cells were rotated for 30 minutes at 4°C and pelleted at 10,000 g for 10 minutes. Pellet was resuspended in 0.4 N Sulphuric Acid and rotated for 30 minutes. Supernatant was collected and 50% (w/v) TCA was added dropwise to 25% final TCA concentration and incubated on ice for 30 minutes before pelleting at 16,000 g for 10 minutes. Pellets were washed three times with ice-cold acetone, allowed to air dry and histones were dissolved in water. 1 µg of purified histones were run on 18% all tris gel using western blot protocol.

#### Reverse Transcription

1 μg of RNA in nuclease free water, up to 16 μL total, were denatured at 65°C for 10 minutes and immediately cooled on ice to chill. cDNA was synthesized using an Eppendorf Mastercycler pro thermal cycler from denatured total RNA by reverse transcription using a 20 μL reaction mixture containing 1 μg RNA, 4 μL 5X iScript Reverse Transcription Supermix for RT-qPCR (BIO-RAD) following the manufacturers protocol.

#### Quantitative PCR (qPCR) Analysis

qPCRs were performed using a BIO-RAD CFX384 Real-Time System in 96 well plates. Reactions were performed using 10 μL reaction mixture containing either 10 ng of DNA or 0.5 μL cDNA, 0.3 μL of 10 μM primer set mixture, 5 μL of iTaq Universal SYBR Green Supermix (BIO-RAD) and nuclease free water. Samples were run using the following program: 95°C for 3 minutes, 45 cycles (increased to 70 cycles for ChIP-qPCR samples) of 5 seconds at 95°C and 20 seconds at 60°C, and finally a melt curve was run from 65 – 95°C raising temperature 0.5 degrees per minute. Samples were run in triplicate. Cycle values were used to determine fold change and normalized to expression of β-Actin or Hdac1.

#### Flow Cytometry

For DNA content analysis 10^6^ cells were washed in PBS (NaCl 137 mM, KCl 2.7 mM, Na2HPO4 10 mM, KH2PO4 1.8 mM) and fixed in ice cold 70% ethanol overnight. Fixed cells were washed and resuspended in PBS containing 0.05% NP40 and incubated RT for 30 minutes. Cells were again washed with PBS and then stained for total DNA content using 10 μg/mL propidium iodide (PI) and 0.2 mg/ml RNase A for 60 minutes before analysis on a FACScan instrument. Data was analyzed using FlowJo (Version 10) with cells gated to remove doublets.

#### Rerep-seq

Rerep-seq was performed as described in ^39^. Briefly, In an 8 strip PCR tube 1 – 5 ug high molecular weight genomic DNA was mixed with 2.5uL 10X Hoechst 33258 (0.1mg/ml) and

2.5uL 10X NEB 4 (50 mM Potassium Acetate, 20 mM Tris-acetate, 10 mM Magnesium Acetate, 1 mM DTT pH 7.9@25°C) to a final volume of 24uL; open tubes were then placed upright in PCR tube rack, covered with a glass plate (3" x 3" glass plate from VWR Vertical Gel Electrophoresis Systems), exposed to 7.5 minutes of glass plate filtered (UVA only) from Stratalinker followed by addition of 0.5uL UDG (5 units of Uracil-DNA Glycosylase), 0.5uL APE1 (10 units of human apurinic/apyrimidinic endonuclease 1) and incubated for 2 hours at 37°C. The digested DNA was repaired with NEB’s FFPE DNA Repair Mix for 30 minutes; mixed with 10X loading dye then run on a 0.8% agarose gel for 15 minutes. The fragmented DNA ranging from 0.1Kb to 3Kb was excised from the gel andextracted with Wizard® SV Gel and PCR Clean-Up System, finally the purified DNA was eluted in 50uL of nuclease free water. Sequencing libraries were constructed using the Illumina Nextera DNA Flex Library Prep Kit (catalog number 20018704) following the manufacturer’s protocol and using 14 cycles of amplification for final library construction. Libraries were constructed using dual index barcodes from the Nextera DNA CD Indexes (catalog number 20018707). Pooled libraries were sequenced by Novogene (Novogene Corporation INC Chula Vista CA) to obtain approximately 10 million reads per yeast sample and 30 million reads per human sample.

#### Western Blot

Whole cell lysates were collected in RIPA buffer (50 mM Tris pH 7.4, 150 mM NaCl, 1% NP-40, 0.25% Sodium Deoxycholate, 1 mM EDTA pH 8.0, 20% glycerol) unless otherwise stated. Samples were sonicated for 10 minutes using 30 second on/off cycle repeats at 70% intensity (using a Qsonica Q800R). 20-50 μg of protein were loaded into all tris gels (6% to 18% depending on target protein sizes) and run at 0.1 amps until the loading buffer dye ran through the gel. Gels were then transferred to BioTrace NT nitrocellulose paper (Pall Corporation #66485) for 200 minutes at 200 mAmps at 4°C. Nitrocellulose paper was stained in Ponceau S staining solution (0.1%(w/v) Ponceau S [Amresco 0860] in 5% acetic acid) to locate protein bands. Ponceau S staining solution was washed off in water and nitrocellulose paper was blocked for 1 hour in blocking solution (5% BSA in PBST). Primary antibody (antibody in PBST containing 5% BSA) was applied with shaking overnight at 4°C. Nitrocellulose was washed 3 times for 5 minutes in PBST and secondary (antibody in PBST containing 5% milk) was applied with shaking at RT for 1 hour. The Nitrocellulose paper was washed 3 times for 5 minutes in PBST and chemiluminescent substrate (SuperSignal West Duro Extended Duration Substrate Thermo Scientific #34075) was applied. Blots were imaged using a LI-COR Odyssey FC instrument running Image Studio software (version 5.2).

#### Cell Survival Assay

Cells were plated in 96-well plates and treated as indicated for 72 hours before performing MTT survival assay using Cell Proliferation Kit 1 MTT (Roche) following the manufacturer’s instructions. Absorbance at 550 nM on was measured on a Biotech Synergy 3 spectrophotometer with Gen5 software version 1.11. Absorbance values were normalized to control sample absorbance (set as maximum) and lowest absorbance (set to zero survival).

#### Immunoprecipitation and Chromatin Immunoprecipitation

Cells were crosslinked by adding 1% formaldehyde to media for 10 minutes at 37°C. Crosslinking was stopped by addition of Glycine to 0.125M and incubating for 5 minutes at room temperature. Cells were washed with PBS and collected and centrifuged. Cells were resuspended in cellular lysis buffer (5mM PIPES pH 8, 85 mM KCl, 0.5% NP40), incubated on ice for 5 minutes and centrifuged. Samples were resuspended in nuclear lysis buffer (50 mM Tris pH 8, 10 mM EDTA pH 8, 1% SDS) and sonicated for 45 minutes using 30 second on/off cycle repeats at 70% intensity (using a Qsonica Q800R). 10 μg crosslinked DNA and 2 μg crosslinked Drosophila control spike-in chromatin was used per immunoprecipitation. Chromatin was pulled down overnight using Invitrogen Protein A Dynabeads with antibodies for H3 (Cell Signaling Technology), H3K27me3 (Abcam) or IgG (Cell Signaling Technology). Beads were washed twice with IP buffer (16.7 mM Tris pH 8, 1.2 mM EDTA pH 8, 167 mM NaCl, 0.1% SDS, 1.84% Triton X100), once with TSE buffer (20 mM Tris pH 8, 2 mM EDTA pH 8, 500 mM NaCl, 1% Triton X100, 0.1% SDS) and LiCl buffer (100mM Tris pH 8, 500 mM LiCl, 1% deoxycholic acid, 1% NP40), and twice with TE (10mM Tris pH 8, 1 mM EDTA pH 8). DNA was eluted from beads with proteinase K and purified with Promega Wizard SV PCR clean up kit. DNA was analyzed with qPCR and compared to 10% input collected prior to pulldown.

#### ChIP-seq Library Preparation, Sequencing and Data Processing

ChIP-seq libraries were prepared using the Illumina NEBNext Ultra II ChIP-Seq sample kit, according to the manufacturer’s protocol. Libraries were validated using the Agilent Technologies 2100 Bioanalyzer. Libraries were sequenced by the University of Colorado Cancer Center Genomics and Microarray Core on Illumina NovaSEQ6000 sequencer as paired end 151x8x8x151. ChIP-seq experiments were aligned to a custom genome combining human hg38 and drosophila dr3 genomes using Bowtie2^85^ (v 2.3.4.1). Sam files were transformed into bam files and sorted using samtools^86^ (v 1.8). Reads aligned to chr1-22 and chrX were isolated and an equal number of reads were randomly selected from each biological replicate for H3K27me3 and Input then sorted, merged and indexed using samtools (version 1.8) and alignment coverage bigwig files were created with deeptools^87^ (v 3.1.3) bamCoverage using CPM with 50 bp binning. Genome-wide and MT locus pearson correlations were determined for H3K27me3 ChIP bigwig coverage files using deeptools^87^ (v 3.1.3) multiBigwigSummary to analyze CPM genome wide and create a multiBigwigSummary file followed by deeptools plotCorrelation to analyze the multiBigwigSummary file and perform correlation analysis. MultiBigwigSummary was run for chromosomes 1-22 and chromosome X with parameters: bins -bs=50. PlotCorrelation was run with parameters: –corMethod pearson –whatToPlot scatterplot. We have provided the FASTA files, subsampled BAM files and averaged bigwig files at our github (https://github.com/Black-Lab-UCDenver/MTDNARereplication). Sequencing files are also deposited in the GEO archive under accession number: GSE179822.

#### ChIP-seq H3K27me3 Peak Calling

ChIP-seq peak calling was performed using Epic2^88^ v 0.0.50) using default parameters with merged randomly subsampled bam files (see ChIP-seq Preparation, Sequencing and Data Processing) as --treatment (H3K27me3) and --control (Input) using –genome hg38.

### CUT&RUN

CUT&RUN experiments were carried out following Epicypher CUT&RUN protocol (version 1.6, August 2020) with minor modifications. Briefly, nuclei from 5x10^5^ cells were isolated with wash buffer (20 mM HEPES-KOH, pH 7.5, 150 mM NaCl, 0.5 mM Spermidine, and 1x protease inhibitor cocktails from Sigma), captured with Concanavalin A conjugated paramagnetic beads (Bangs Laboratory Inc.) and incubated while nutating with 0.5 µl primary antibody for EZH2 (Cell Signaling Technology), or SUZ12 (Cell Signaling Technology) in antibody buffer (20 mM HEPES-KOH, pH 7.5, 150 mM NaCl, 0.5 mM Spermidine, 0.02% digitonin, 2 mM EDTA and 1x protease inhibitor cocktails from Sigma) overnight. After washing with digitonin buffer (20 mM HEPES-KOH, pH 7.5, 150 mM NaCl, 0.5 mM Spermidine, 0.02% digitonin and 1x protease inhibitor cocktails from Sigma), pAG-MNase (Epicypher Inc) was added at a 1:20 ratio and incubated for 10 minutes at RT. The nuclei were washed again and placed on ice. To activate pAG-MNase, CaCl2 was added to a final concentration of 2 mM. The reaction was inubated for 2 hours while nutating at 4°C and stopped by addition of STOP buffer (340 mM NaCl, 20 mM EDTA, 4 mM EGTA, 50 mg/mL RNase A and 40 mg/mL glycogen). The protein-DNA complex was released by incubating for 10 minutes at 37°C. DNA was extracted using Qiagen Minelute PCR purification kit. Purified DNA was used for library preparation.

#### CUT&RUN Library Preparation, Sequencing and Data Processing

CUT&RUN libraries were prepared using the Illumina NEBNext Ultra II ChIP-Seq sample kit, according to the manufacturer’s protocol. Libraries were validated using the Agilent Technologies 2100 Bioanalyzer. Libraries were sequenced by the University of Colorado Cancer Center Genomics and Microarray Core on Illumina NovaSEQ6000 sequencer as paired end 151x8x8x151. CUT&RUN experiments were adaptor trimmed using BBtools bbduk (v 38.87) using supplied adaptor reference file adapters.fa with parameters ktrim=r k=23 mink=11 hdist=1 tpe tbo then aligned to a custom genome combining human hg38 and *E. coli* mg1655 genomes using Bowtie2 (v 2.3.4.1). Sam files were transformed into bam files, mapped reads with insert size greater than or equal to 150 bp were isolated, sorted and indexed using samtools (version 1.8). CUT&RUN alignment coverage bigwig files were created with deeptools bamCoverage (v 3.1.3) using CPM with 50 bp binning. Biological replicates were averaged at the bin level using WiggleTools^89^ (v 1.2) using the mean operation and the resulting wig file was converted to bigwig using wigToBigWig command. We have provided the FASTA and averaged bigwig files at our github (https://github.com/Black-Lab-UCDenver/MTDNARereplication). Sequencing files are also deposited in the GEO archive under accession number: GSE179823.

#### Rerep-seq and Whole Genome Sequencing Analysis

Human FASTQ files were aligned using BWA-mem2^86^ (v 2.0, RRID:SCR_010910) to HG38 from UCSC. Sam files were transformed into bam files and blacklisted using samtools (v 1.2, RRID:SCR_002105). Blacklists were the same as previously used for Rerep-seq analysis (Menzel et al., 2020). Bedtools^90^ (v 2.26.0, RRID:SCR_006646) was used to generate bedGraphs and bigWigs, which were binned to 1kb (for whole chromosome visualization and additional analysis), or 50bp (for comparison to CHIP-Seq data), and then normalized by RPM and mitochondrial reads using a custom R script (v3.6, RRID:SCR_001905, (Menzel et al., 2020)). Biological replicate bedGraphs and bigWigs were combined by averaging the signaling per bin and used for further analysis. Yeast FASTQ files were processed in the same fashion as the human files, and aligned to a custom version of UCSC SacCer3, and binned using a 100bp window (used for further analysis) or 50bp (used for chromosome visualization). The CUP1 gene is exactly duplicated in the yeast genome, so we replaced the CUP1.1 gene copy with ambiguous nucleotides “N” to force alignment to CUP1.2 rather than allow for random alignment between the two copies. We have provided the FASTA for this genome, as well as the BWA index at our github (https://github.com/Black-Lab-UCDenver/MTDNARereplication). Sequencing files are also deposited in the GEO archive under accession numbers: GSE165865 (human data) and GSE168566 (yeast data).

#### Structural Variant Analysis

Structural variants were called using GRIDSS (v1.0) ^56^ on default settings. VCF outputs were transformed into bedPe files using the StructuralVariantAnnotation R package (v1.4.0). Double breakpoint variants with a GRIDSS score over 50 for human and over 150 for yeast were selected for further analysis. We identified variants that connect the MT locus to other chromosomal regions. For the purposes of this analysis, we defined the metallothionine locus as chr16:56,520,000-56,720,000 in HG38, and we defined the CUP1 locus was chrVIII:210,500-218,000 in SacCer3. Structural variants that originated from these loci were displayed using R circularize (v0.4.10,).

#### Identification of Homology and Microhomology

To determine if there is homology between the origin (in MT or Cup1 locus) and insertion (other chromosomal location) for the structural variants returned by GRIDSS (v1.0), we used a locally installed version of NCBI BLASTn (v2.6.0, RRID:SCR_001598) to determine if there was alignment (homology) between the origin and insertion sequences.

To evaluate homology, we defined 1 kb segments, centered on the middle of the origin and the insertion for each variant pair (-/+ 500 bp from the GRIDSS predicted breakpoint). Individual BLAST data bases were generated for each origin, and the insertion pair was evaluated using the default parameters for BLASTn with an E-value less than 1. We generated two different controls for this analysis: a random genome control as well as a scrambled control. The scrambled control was generated in R using the stringi package (v1.4.6) to scramble the original insertion sequence. The random genomic sequence was generated by randomly picking a 1kb region from the same genome as the original sample, and then using bedtools (v2.26.0, RRID:SCR_006646) to retrieve the FASTA sequences. The random genomic loci and the scrambled sequences were also aligned to the origin and insertion sequences. For all analyses, the sequence in the 1 kb segment with the best E-value was chosen. For sequences with multiple alignments with the same E-value, the longest homologous stretch was chosen to break the tie.

To evaluate microhomology, we defined 50bp segments, centered on the middle of the origin and insertion of each variant pair (-/+25 bp from the GRIDSS predicted breakpoint). Individual BLAST databases were created for each origin, and insertion, scrambled control, and random genomic controls were evaluated for microhomology by using an E-value less than 1 and a seed length of four base pairs to allow for the identification of short stretches of homology with a minimum of four base pairs of homology. The random genomic loci and the scrambled sequences were also aligned to the origin and insertion sequences. For all analyses, the sequence in the 1 kb segment with the best E-value was chosen. For sequences with multiple alignments with the same E-value, the longest homologous stretch was chosen to break the tie.

The significance of an E-value is inverse to its numerical magnitude: the smaller the E-value, the more significant the homology. To reflect the magnitude of these values visually, we displayed the data as either 1/E-value or log10(1/E-value). The number of variants with homology between the origin and insertion was displayed as a bar plot using the R package ggplot2 (v3.3.2). Significance was calculated using a chi-squared test using the random genomic insertions or the scrambled insertion as the expected. Boxplots were used to illustrate both the length of homology in base pairs and the degree of homology as reflected by the log10(1/E-value) using the R package ggplot2 (v3.3.2). Using R, a non-parametric Kruskal– Wallis test with a post hoc Wilcoxon–Mann–Whitney test was used to measure significance for the log10(1/E-value) plots as well as the length of homology. Histogram density plots were generated to display the frequency of log10(1/E-value) and average traces of 1/E-value across all origin-insertion variant pairs were generated to display the location of homology with respect to the breakpoint, these figures were generated using ggplot2 (v3.3.2). A Kolmogorov–Smirnov test was used to quantify statistical similarity between the log10(1/E-value) distributions of GRIDSS reported variant pairs and either the scrambled insertions or random genomic insertions. To make the homology trace figure, we created a vector of zeros the length of each variant fragment, and replaced the locations where homology was identified with either the 1/E-value or log10(1/E-value). An average value was generated across all variants by vector position resulting in an average trace of homology location as well as magnitude of significance. The final figure was generated using the R package ggplot2(v3.3.2).

#### Extrachromosomal Circle Analysis

Extrachromosomal DNA circles can be inferred by isolating reverse-forward read pairs from paired sequencing data. We isolated these read pairs using samtools (v1.2, RRID:SCR_002105) and sam flag 81 and created bedPe files using bedtools (v2.26.0, RRID:SCR_006646) for further analysis. These reads were filtered to isolate read pairs originating within or immediately adjacent to the metallothionine locus in human samples and the CUP1 locus in yeast (56,550,000-56,700,000 in human and 211,000-218,000 in yeast). These read pairs were displayed using the packages Gviz^91^ (v 3.11) and GenomicInteractions^92^ (v 1.22).

#### Repeat Element Enrichment Analysis

Repeat element enrichment within structural variants was performed using the DFAM database (V3.2). RepeatMasker (V4.1.1) was used with default parameters to identify repeats from DFAM genome wide, returning 5,607,738 repeats. Bedtools intersect (v2.26.0, RRID:SCR_006646) was used with default parameters, which looks for one base pair or more of overlap between two features, to identify repeats and homologous SVs that occurred at the same locations. R was used perform a fisher exact test to determine repeat sub-type as well as family enrichment, using the genome wide distribution of sub-type or family of repeats as the control compared to the distribution of repeats among our homologous SVs.

### QUANTIFICATION AND STATISTICAL ANALYSIS

#### Statistical Analysis

All pairwise comparisons were done using two-tailed Student’s t -test unless otherwise stated. Significance was determined if the P-value was < 0.05. All FISH experiments were carried out with at least two independent siRNAs in at least two independent transfections and at least 150 nuclei per replicate were counted for all FISH studies conducted. Error bars represent S.E.M.

## DATA AND CODE AVAILABILITY

The data analyzed in this study can be found at the GEO archive (accession numbers: GSE numbers: GSE165865, GSE165866, GSE179822 and GSE179823). All code used in this study can be found on our github page: (https://github.com/Black-Lab-UCDenver/MTDNARereplication). All supplementary files such as blacklists, scrambled and random genomic controls for the homology analysis, and the custom saccer3 genome can all be found on our github page: (https://github.com/Black-Lab-UCDenver/MTDNARereplication).

## DECLARATION OF INTERESTS

The authors declare no competing interests.

---

Tableof Important Resources

Table of Important Resources
| REAGENT or RESOURCE | SOURCE | IDENTIFIER |
| --- | --- | --- |
| **Antibodies** |  |  |
| Rabbit monoclonal anti-H3 – ChIP formulated | Cell Signaling | 4620S;<br>RRID:AB_1904005 |
| Rabbit monoclonal anti-H3 | Cell Signaling | 4499S;<br>RRID:AB_10544537 |
| Rabbit-monoclonal anti-H3K27me3 | abcam | Ab192985;<br>RRID:AB_2650559 |
| Mouse monoclonal anti-Chk1 | Santa cruz biotechnology | SC8408;<br>RRID:AB_627257 |
| Mouse-monoclonal anti-Actinin | Santa cruz biotechnology | SC17829;<br>RRID:AB_626633 |
| Normal Rabbit IgG | Cell Signaling | 2729S, 2729S, RRID:AB_1031062 |
| Rabbit monoclonal anti-EZH2 | Cell Signaling | 5246S,<br>RRID:AB_10694683 |
| Rabbit monoclonal anti-SUZ12 | Cell Signaling | 3737S,<br>RRID:AB_2196850 |
| **Chemicals, Peptides, and Recombinant Proteins** |  |  |
| Sodium Citrate | Amresco | 0101-1KG |
| Pepsin | VWR | 0719-25G |
| Dapi | Sigma | D9542 |
| Trichloroacetic Acid | VWR | 0133-500G |
| Prolong Gold antifade reagent | Invitrogen | P36934 |
| 1,4 Dithiotheritol | Sigma | 10708984001 |
| Manganese(II) Chloride Solution | Sigma | M1787 |
| Sodium Deoxycholate | VWR | 0613-100G |
| Lithium Chloride | J.T. Baker | 2370-07 |
| Deoxychloric Acid Sodium Salt | VWR | 0613-100G |
| Dynabeads protein A | Invitrogen | 10002D |
| Concanavalin A Beads | Bangs Laboratories Inc | BP531 |
| pAG-Mnase | Epiccypher | 15-1016 |
| Digitonin High Purity | Millipore | 300410-250MG |
| cOmplete min EDTA free protease inhibitor cocktail | Roche Diagnostics | 11836170001 |
| Nitrocellulose paper | Pall Corporation | 66485 |
| Ponceau S | Amresco | #0860 |
| Hydroxy Urea | Sigma | H8627-5G |
| UNC1999 | Selleckchem | S7165 |
| Cadmium Sulphate | Arcos Organics | 270565000 |
| Cisplatin | Selleckchem | S1166 |
| VE821 | Selleckchem | S8007 |
| PTC209 | Selleckchem | S7372 |
| PRT4165 | Selleckchem | S5315 |
| EX527 | Selleckchem | S1541 |
| Sirtinol | Selleckchem | S2804 |
| CP2 | Selleckchem | S8601 |
| CPI455 | Selleckchem | S8287 |
| **Critical Commercial Assays** |  |  |
| miRNeasy Minikit | Qiagen | 1038703 |
| Rnase-Free Dnase set | Qiagen | 79254 |
| QIASol Lysis Reagent | Qiagen | 1023537 |
| SuperSignal West Duro Extended Duration Substrate | Thermo Scientific | 34075 |
| NEBNext Ultra II DNA Library Prep Kit for Illumina | NEB | E7645L |
| NEBNext® Multiplex Oligos for Illumina® (Index Primers Set 1) | NEB | E7335L |
| QIAgen MinElute PCR Purification Kit | Qiagen | 28004 |
| **Deposited Data** |  |  |
| Raw and analyzed data – yeast rerep-seq | This paper | GEO: GSE168566 |
| Raw and analyzed data – human rerep-seq | This paper | GEO: GSE165865 |
| Raw and analyzed data – human ChIP-seq | This paper | GEO: GSE179822 |
| Raw and analyzed data – human CUT&RUN | This paper | GEO: GSE179823 |
| **Experimental Models: Cell Lines** |  |  |
| Human: MDA-MB-231 | ATCC | HTB-26 |
| Human: RPE | Dyson Lab | Black et al., 2013 |
| **Experimental Models: Organisms/Strains** |  |  |
| *S. cerevisiae* strain YJB1 | Sclafani Laboratory | Zhong et. al., 2013 |
| *S. cerevisiae* strain YJB21 *cup1Δrsc30Δ* | This paper | N/A |
| *S. cerevisiae* strain YJB26 CUP1S | This paper | N/A |
| *S. cerevisiae* strain YJB30 *cup1Δ* | This paper | N/A |
| *S. cerevisiae* strain YJB31 Cup2b | This paper | N/A |
| *S. cerevisiae* strain YJB36 Cup2b2 | This paper | N/A |
| *S. cerevisiae* strain YJB33 pGAL-CUP1S | This paper | N/A |
| *S. cerevisiae* strain YJB41 ARS1-pGAL-CUP1S | This paper | N/A |
| **Oligonucleotides** |  |  |
| See table S1 for primers |  |  |
| **Recombinant DNA** |  |  |
| 16qhet FISH probe | Cytocell | MPD6562 |
| 16qter Subtelomere Specific FISH Probe | Cytocell | LPT16QG |
| MT in Texas Red FISH Probe | Cytocell | MPD6561 |
| Chromosome 1 Classic Satellite FISH Probe | Cytocell | LPE001G-A |
| Chromosome 8 Alpha Satellite FISH Probe | Cytocell | LPE008R |
| Silencer Select Negative Control siRNA | Life Technologies | s813 |
| EZH2 siRNA | Life Technologies | s4916 |
| EZH2 siRNA | Life Technologies | s4918 |
| SUZ12 siRNA | Life Technologies | s23967 |
| SUZ12 siRNA | Life Technologies | s23969 |
| EED siRNA | Life Technologies | s16626 |
| EED siRNA | Life Technologies | s16624 |
| **Software and Algorithms** |  |  |
| Rstudio v 1.0.153 | RStudio | https://rstudio.com |
| Bowtie2 | Langmead and Salzberg, 2012 | http://bowtie-bio.sourceforge.net/bowtie2/index.shtml |
| Samtools | Li et al., 2009 | http://www.htslib.org/ |
| deeptools | Ramírez et al., 2016 | https://deeptools.readthedocs.io/en/develop/ |
| wiggletools | Zerbino et al., 2014 | https://github.com/Ensembl/WiggleTools |
| BWA-mem2 | BWA-mem2 | https://github.com/bwa-mem2/bwa-mem2 |
| bedtools | Quinlan and Hall, 2010 | https://bedtools.readthedocs.io/en/latest/ |
| GRIDSS | Cameron et al, 2017 | https://github.com/PapenfussLab/gridss |
| NCBI BLASTn | NCBI, 2018 | https://blast.ncbi.nlm.nih.gov/Blast.cgi |
| Gviz | Helaers et al, 2011 | https://www.bioconductor.org/packages/release/bioc/html/Gviz.html |
| GenomicInteractions | Harmston et al., 2015 | https://www.bioconductor.org/packages/release/bioc/html/GenomicInteractions.html |
| Epic2 | Stovner et al, 2019 | https://github.com/biocore-ntnu/epic2 |

## Notes

### Competing Interest Statement

The authors have declared no competing interest.

