## Supplemental Figures and Legends for "Transition from Transient DNA Rereplication to Inherited Gene Amplification Following Prolonged Environmental Stress"

### Figure S1

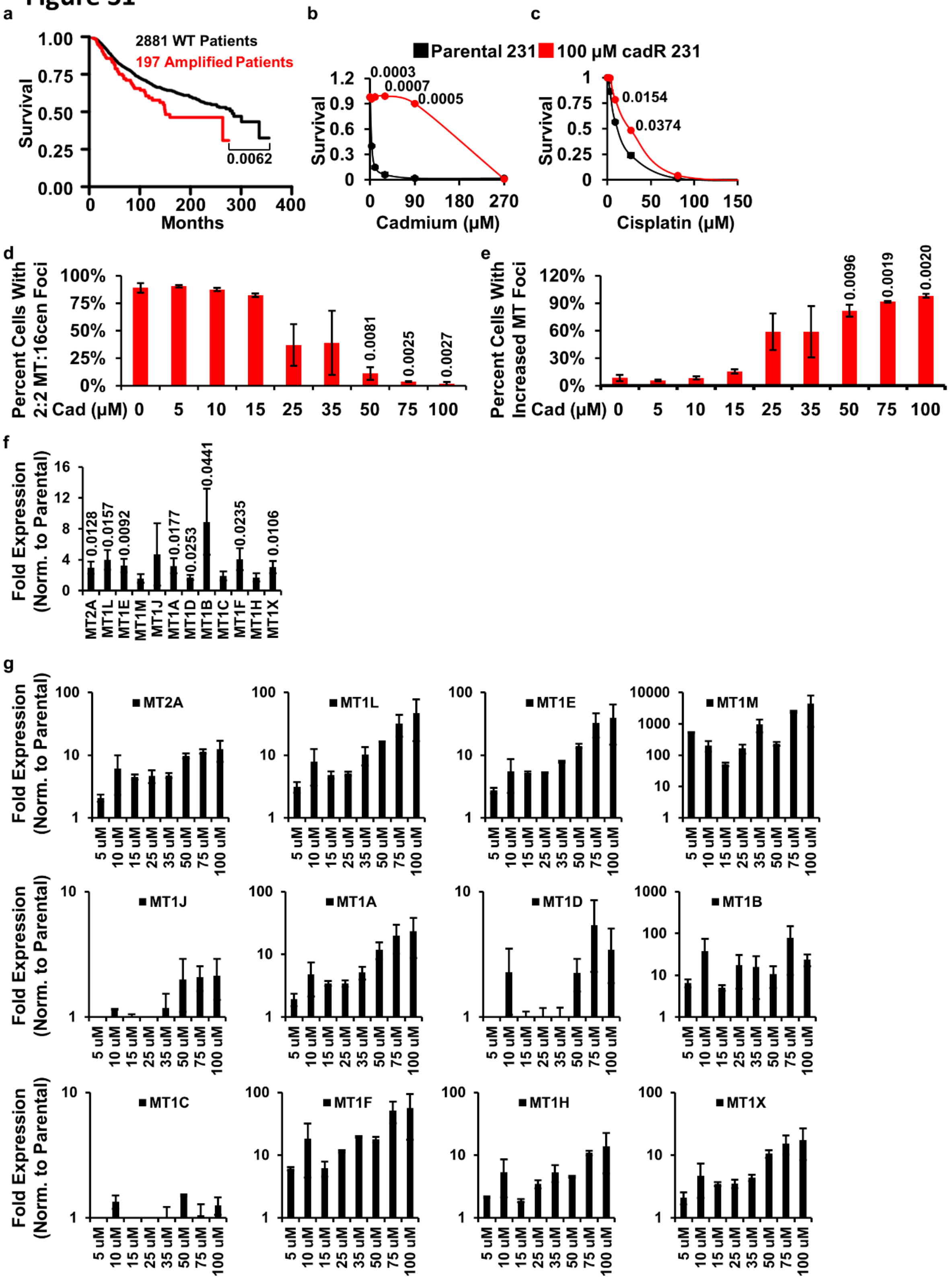

a
2881 WT Patients
197 Amplified Patients
Survival
Months
0.0062
b
c
Parental 231
100 µM cadR 231
0.0003
0.0007
0.0005
Survival
Cadmium (µM)
0.0154
0.0374
Survival
Cisplatin (µM)
d
Percent Cells With 2:2 MT:16cen Foci
Cad (µM)
0.0081
0.0025
0.0027
e
Percent Cells With Increased MT Foci
Cad (µM)
0.0096
0.0019
0.0020
f
Fold Expression (Norm. to Parental)
0.0128
0.0157
0.0092
0.0177
0.0253
0.0441
0.0235
0.0106
MT2A MT1L MT1E MT1M MT1J MT1A MT1D MT1B MT1C MT1F MT1H MT1X
g
Fold Expression (Norm. to Parental)
MT2A
MT1L
MT1E
MT1M
MT1J
MT1A
MT1D
MT1B
MT1C
MT1F
MT1H
MT1X
5 uM 10 uM 15 uM 25 uM 35 uM 50 uM 75 uM 100 uM


19

**Figure S1: Related to Figure 1. Cells Develop Progressive MT Amplifications in Response to Cadmium Exposure**

(a) Kaplan Meyer survival curve for TCGA and METABRIC breast cancer patients separated by Metallothionein amplification (red) and non-amplified (black). (b) MDA-MB-231CadR cells survive higher cadmium concentrations as the cells develop cadmium resistance. (n=2 independent cell cultures). (c) MDA-MB-231CadR cells survive higher cisplatin concentrations as the cells develop cadmium resistance. (n=2 independent cell cultures). (d) MDA-MB-231CadR cells have lower percent of cells with 2:2 (MT:16cen) foci count by DNA-FISH as cadmium concentration increases. (n=2 biological replicates). (e) MDA-MB-231CadR cells have a higher percent of cells with increased count of MT foci by DNA-FISH as cadmium concentration increases. (n=2 biological replicates). (f) Acute 72hr cadmium treatment in MDA-MB-231 cells induces expression of MT genes. (n=4 independent cell cultures). (g) MDA-MB-231CadR cells have increased expression of MT genes as the cells develop cadmium resistance. (n=2 biological replicates). Note MT4, MT3 and MT1G exhibit little or no expression in these cells and are not depicted. Error bars represent the S.E.M. * indicates $p < 0.05$ by two-tailed Student's t-test.

### Figure S2

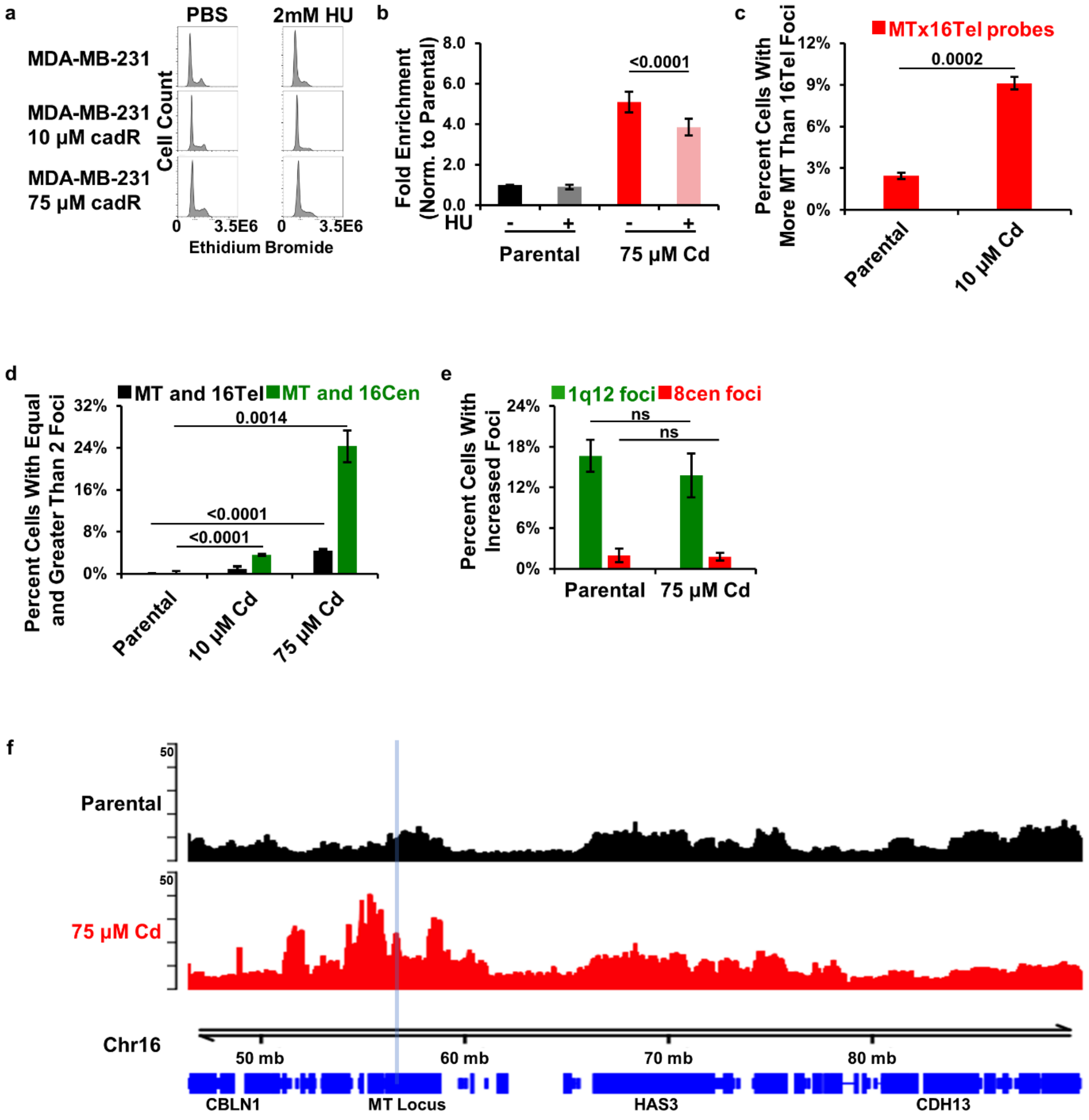
a
PBS
2mM HU
MDA-MB-231
MDA-MB-231 10 µM cadR
MDA-MB-231 75 µM cadR
Cell Count
0 3.5E6 0 3.5E6
Ethidium Bromide
b
Fold Enrichment (Norm. to Parental)
8.0
6.0
4.0
2.0
0.0
HU
- + - +
Parental
75 µM Cd
<0.0001
c
MTx16Tel probes
Percent Cells With More MT Than 16Tel Foci
12%
9%
6%
3%
0%
0.0002
Parental
10 µM Cd
d
MT and 16Tel
MT and 16Cen
Percent Cells With Equal and Greater Than 2 Foci
32%
24%
16%
8%
0%
0.0014
<0.0001
<0.0001
Parental
10 µM Cd
75 µM Cd
e
1q12 foci
8cen foci
Percent Cells With Increased Foci
24%
18%
12%
6%
0%
ns
ns
Parental
75 µM Cd
f
Parental
75 µM Cd
50
50
Chr16
50 mb
60 mb
70 mb
80 mb
CBLN1
MT Locus
HAS3
CDH13

36

**Figure S2: Related to Figure 2. Cadmium Induces Transient Site-Specific Gains of the MT Locus**

(a) MDA-MB-231 parental and cadmium cells treated for 20 hours with HU have decreased proportion of cells in the G2 and M phases of the cell cycle. (b) 75 µM MDA-MB-231 cells have reduced MT locus by qPCR when arrested for 20 hours with HU. (n=3 independent cell cultures). (c) MDA-MB-231CadR cells treated with 10 µM cadmium have increased percent of cells with more MT than chromosome 16 telomere (16-qtel) foci by DNA-FISH. (n=3 independent cell cultures). (d) MDA-MB-231CadR cells treated with 10 µM or 75 µM cadmium have significantly less MT and 16-qtel gains than MT and 16cen gains. (n=3 independent cell cultures). (e) MDA-MB-231CadR cells do not exhibit TSSG of 1q12 or the chromosome 8 centromere. (n=3 independent cell cultures). (f) Rerep-seq analysis of 75 µM CadR cells shows that rereplication occurs throughout the MT locus, the area surrounding the MT locus and extends towards the chromosome 16 centromere. Tracks represent the average of two biological replicates. Error bars represent the S.E.M. P-values determined using two-tailed Student's t-test.

#### Figure S3

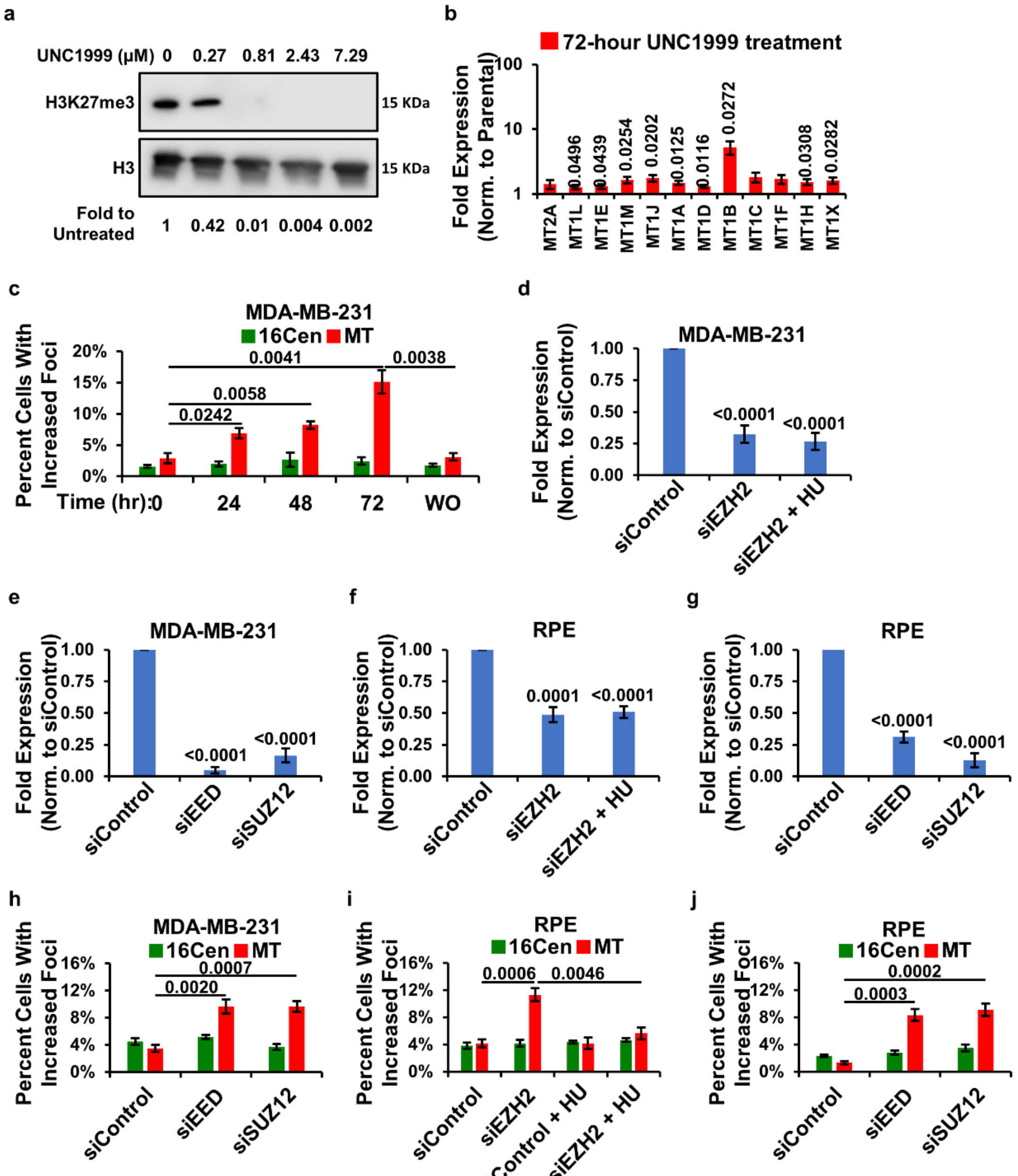
a
UNC1999 (μM) 0 0.27 0.81 2.43 7.29
H3K27me3
15 KDa
H3
15 KDa
Fold to Untreated 1 0.42 0.01 0.004 0.002
b
72-hour UNC1999 treatment
Fold Expression (Norm. to Parental)
100
10
1
0.0496
0.0439
0.0254
0.0202
0.0125
0.0116
0.0272
0.0308
0.0282
MT2A MT1L MT1E MT1M MT1J MT1A MT1D MT1B MT1C MT1F MT1H MT1X
c
MDA-MB-231
16Cen MT
Percent Cells With Increased Foci
20% 15% 10% 5% 0%
0.0041
0.0038
0.0058
0.0242
Time (hr):0 24 48 72 WO
d
MDA-MB-231
Fold Expression (Norm. to siControl)
1.00 0.75 0.50 0.25 0.00
<0.0001
<0.0001
siControl siEZH2 siEZH2 + HU
e
MDA-MB-231
Fold Expression (Norm. to siControl)
1.00 0.75 0.50 0.25 0.00
<0.0001
<0.0001
siControl siEED siSUZ12
f
RPE
Fold Expression (Norm. to siControl)
1.00 0.75 0.50 0.25 0.00
0.0001
<0.0001
siControl siEZH2 siEZH2 + HU
g
RPE
Fold Expression (Norm. to siControl)
1.00 0.75 0.50 0.25 0.00
<0.0001
<0.0001
siControl siEED siSUZ12
h
MDA-MB-231
16Cen MT
Percent Cells With Increased Foci
16% 12% 8% 4% 0%
0.0007
0.0020
siControl siEED siSUZ12
i
RPE
16Cen MT
Percent Cells With Increased Foci
16% 12% 8% 4% 0%
0.0006
0.0046
siControl siEZH2 siControl + HU siEZH2 + HU
j
RPE
16Cen MT
Percent Cells With Increased Foci
16% 12% 8% 4% 0%
0.0002
0.0003
siControl siEED siSUZ12

52

### Figure S3

k

| Treatment | Parental | 10 µM Cd |
|---|---|---|
| Region | Genome | Genome |
| ChIP Reads | 122489484 | 122489484 |
| Peak Count | 57337 | 52310 |
| Avg Peak Score | 77.353 | 76.498 |
| Std Dev | 35.307 | 36.534 |

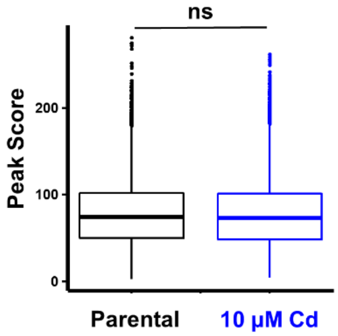


l

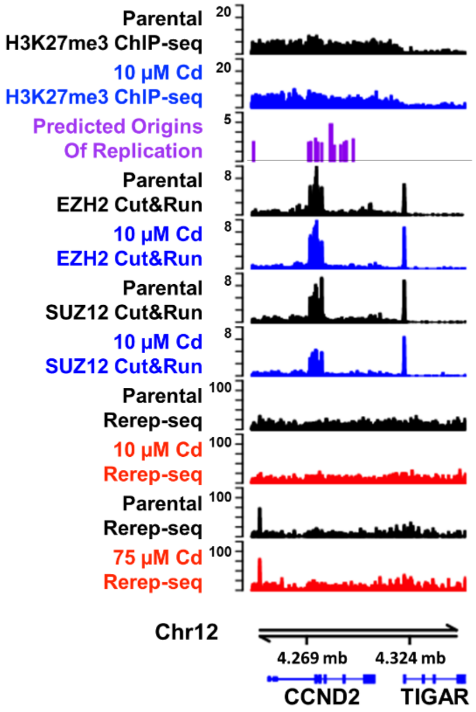


53

**Figure S3: Related to Figure 3. EZH2 and H3K27me3 Suppress TSSG of the MT Locus**

(a) Western blot of acid extracted histones. UNC1999 decreases H3K27me3 levels of MDA-MB-231 cells in a dose dependent manor. (n=4 independent treatments). (b) Acute treatment of MDA-MB-231 cells with 810 nM UNC1999 induces expression of MT genes. (n=3 independent treatments). (c) Acute treatment (72 hours) of MDA-MB-231 cells with 810 nM UNC1999 results in a significant increase in the percent of cells with increased MT foci by DNA-FISH for treatments between 24-72 hours which is lost 24 hours after removal of UNC1999. (n=3 independent treatments). (d-e) Expression of EZH2, EED and SUZ12 following depletion with the indicated siRNA in MDA-MB-231 cells. (n=4, two independent transfections of two different siRNA). (f-g) Expression of EZH2, EED and SUZ12 following depletion with the indicated siRNA in RPE cells. (n=4, two independent transfections of two different siRNA). (h) Treatment of MDA-MB-231 cells with siRNA targeting EED and SUZ12 produce focal gains of the MT locus through DNA-FISH. (n=4, two independent transfections of two different siRNA). (i) Treatment of RPE cells with siRNA targeting EZH2 produce focal gains of MT locus through DNA-FISH, which are lost when cells are arrested in G1/S with HU. (n=4, two independent transfections of two different siRNA). (j) Treatment of RPE cells with siRNA targeting EED and SUZ12 produce focal gains of the MT locus through DNA-FISH. (n=4, two independent transfections of two different siRNA). (k) Broad H3K27me3 peaks called by Epic2 for parental and 10 µM Cd MDA-MB-231 cells show no significant difference in distribution of peak scores genome wide. (l) H3K27me3 levels, EZH2 and SUZ12 localization and Rerep-seq at the CCND2 gene locus in parental and 10 µM Cd MDA-MB-231 cells. H3K27me3 ChIP-seq tracks represent the mean of 3 biological replicates. EZH2 and SUZ12 CUT&RUN and Rerep-seq tracks represent the mean of 2 biological replicates. Error bars represent the S.E.M. P-values determined using two-tailed Student's t-test, box plot in figure (k) utilizes Bonferroni correction for large number of values.

#### Figure S4

a

| | Diploid | Integrated Inherited Amplification |
|---|---|---|
| | 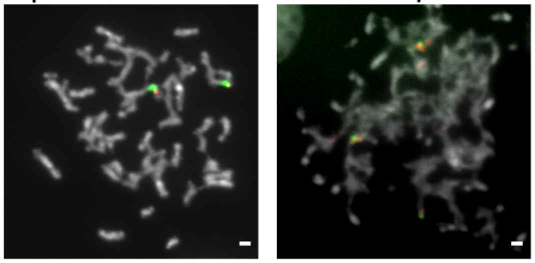 | |
| DMSO | 98.18%<br>54 of 55 | 1.82%<br>1 of 55 |
| UNC1999 | 38%<br>19 of 50 | 62%<br>31 of 50 |

b

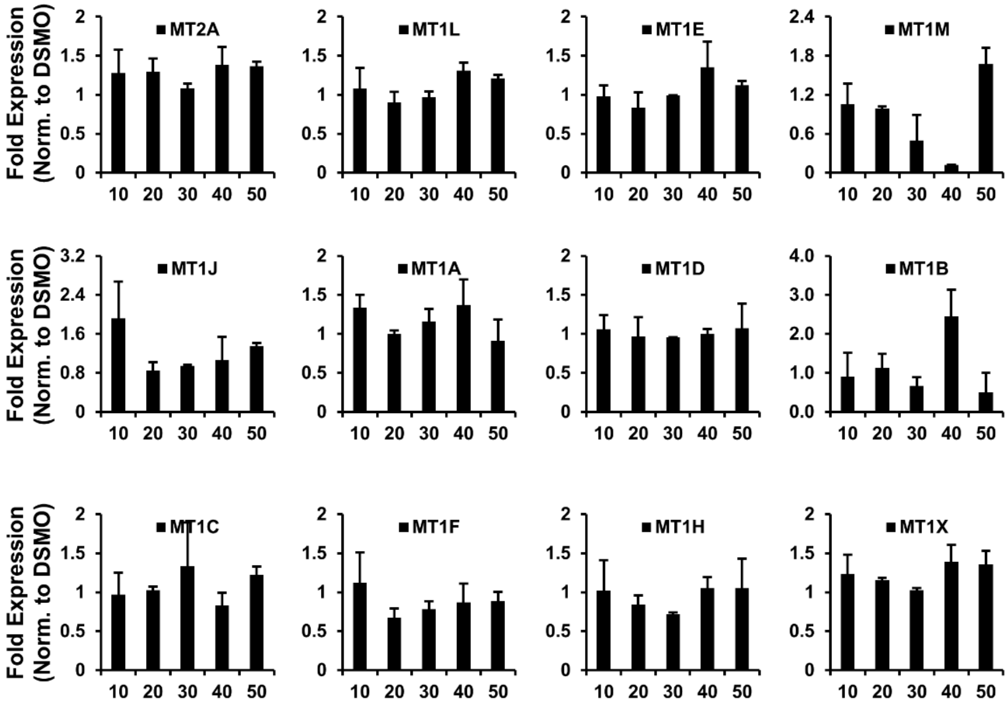


78

**Figure S4**

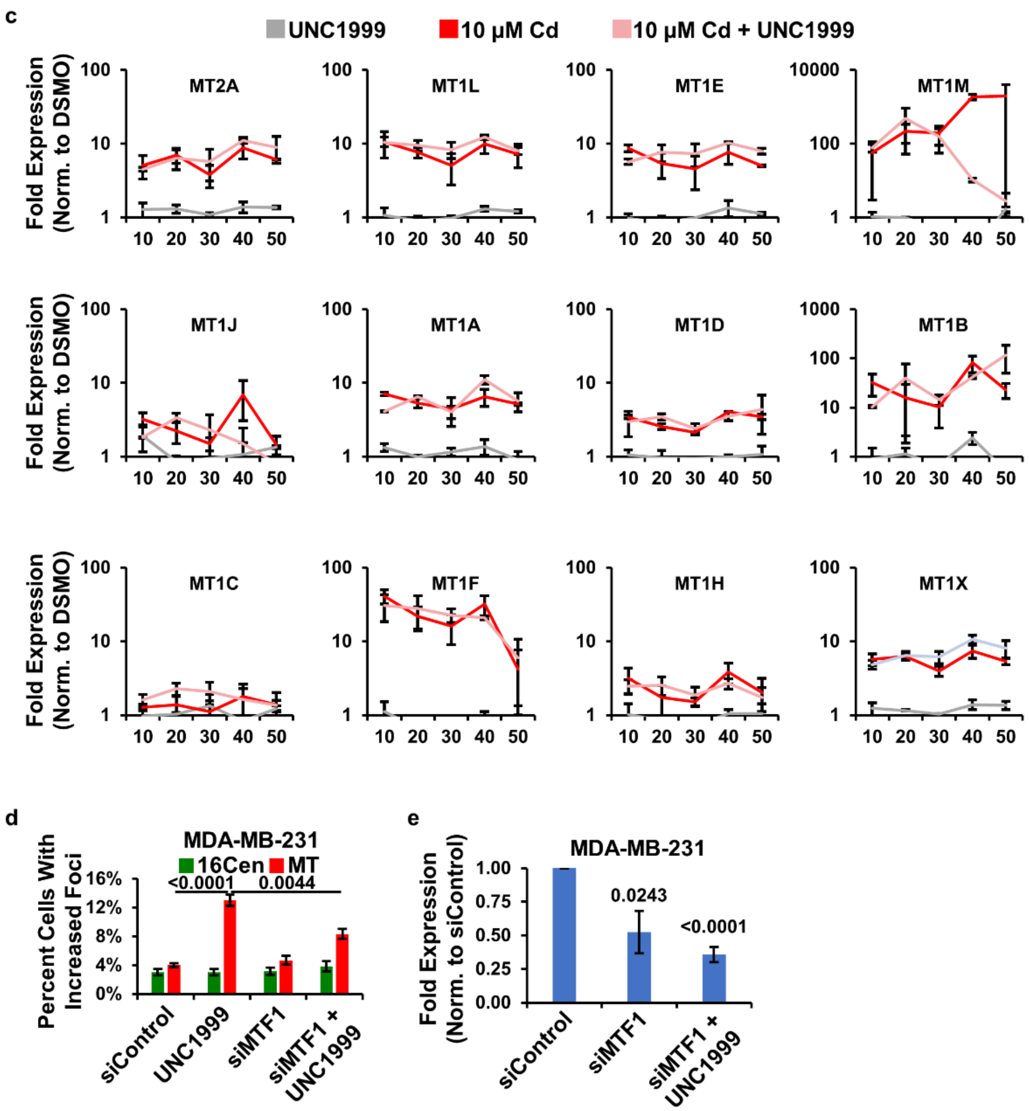
c
UNC1999
10 µM Cd
10 µM Cd + UNC1999
Fold Expression (Norm. to DSMO)
MT2A
MT1L
MT1E
MT1M
MT1J
MT1A
MT1D
MT1B
MT1C
MT1F
MT1H
MT1X
10 20 30 40 50
d
MDA-MB-231
16Cen
MT
Percent Cells With Increased Foci
<0.0001
0.0044
16%
12%
8%
4%
0%
siControl
UNC1999
siMTF1
siMTF1 + UNC1999
e
MDA-MB-231
Fold Expression (Norm. to siControl)
1.00
0.75
0.50
0.25
0.00
0.0243
<0.0001
siControl
siMTF1
siMTF1 + UNC1999

79

**Figure S4: Related to Figure 4. Prolonged EZH2 Inhibition Results in Inherited Metallothionein Amplification Without the Selective Pressure of Cadmium**

(a) Metaphase spread with DNA-FISH for MDA-MB-231 cells following prolonged treatment with 810 nM UNC1999 display increase population of cells with tandem duplications, MT gains without chromosome 16 centromere and repeat tandem duplications of the MT locus. Numbers indicate percentage of cells scored that exhibited amplification of the indicated types ($n>50$ as indicated). (b) Prolonged 810 nM UNC1999 treated MDA-MB-231 cells have increased expression of MT genes as the cells develop cadmium resistance. Note MT4, MT3 and MT1G exhibit little or no expression in these cells and are not depicted. (n=2 biological replicates) (c) Prolonged 810 nM UNC1999 treatment, 10 µM Cd and combined 810 nM UNC1999 with 10 µM Cd treated MDA-MB-231 cells have increased expression of MT genes as the cells develop cadmium resistance. Note MT4, MT3 and MT1G exhibit little or no expression in these cells and are not depicted. (n=2 biological replicates) (d) Acute treatment (72 hours) of MDA-MB-231 cells with 810 nM UNC1999 produce focal gains of MT locus through DNA-FISH, which are lost when cells are treated with siRNA targeting MTF1. (n=4, two independent transfections of two different siRNA) (e) Expression of MTF1 following depletion with the indicated siRNA in MDA-MB-231 cells. (n=4, two independent transfections of two different siRNA). Error bars represent the S.E.M. P-values determined using two-tailed Student's t-test.

#### Figure S5

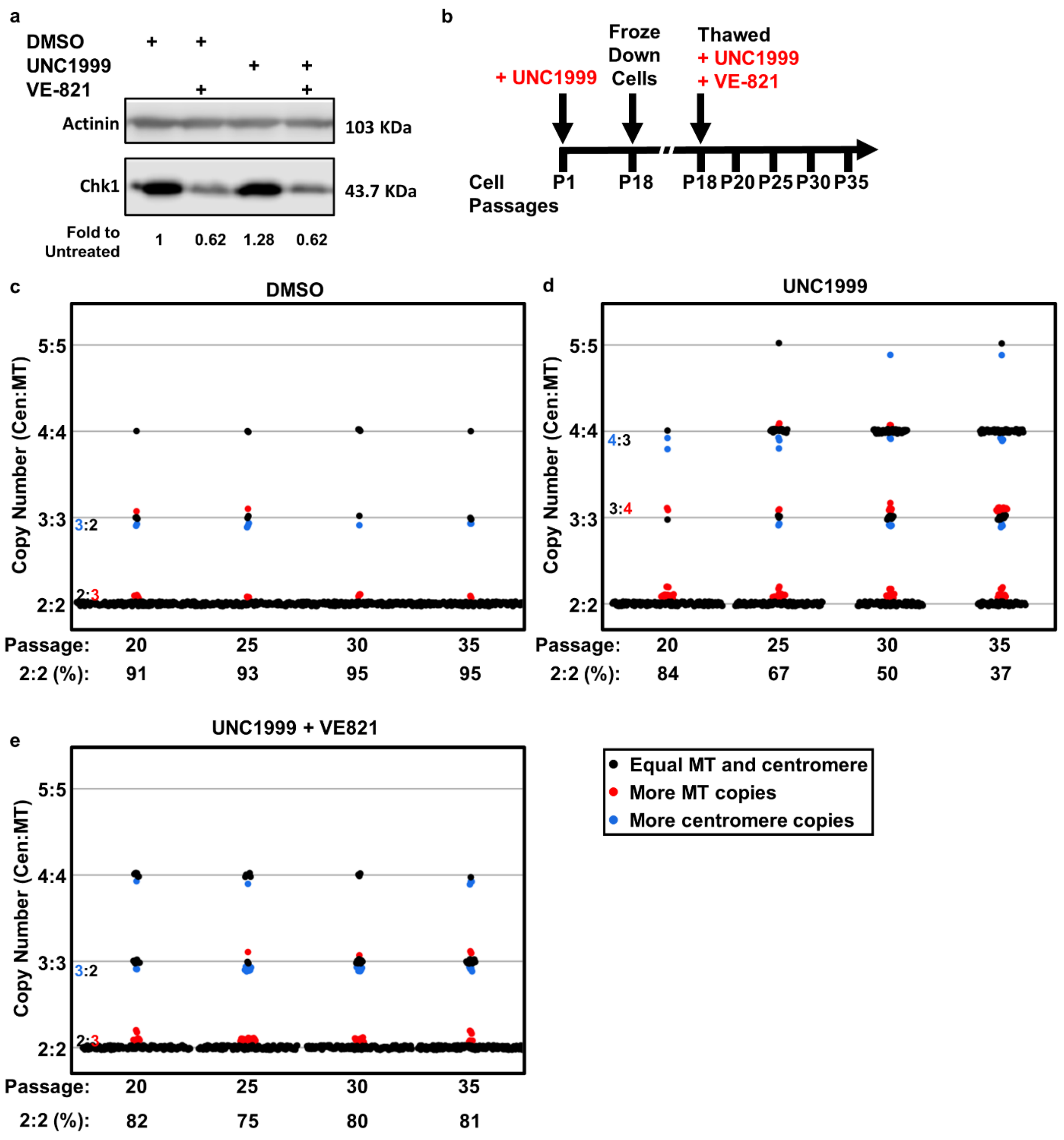

a
DMSO
UNC1999
VE-821
Actinin
103 KDa
Chk1
43.7 KDa
Fold to Untreated
1
0.62
1.28
0.62
b
Froze Down Cells
Thawed
+ UNC1999
+ VE-821
+ UNC1999
Cell Passages
P1
P18
P18
P20
P25
P30
P35
c
DMSO
Copy Number (Cen:MT)
5:5
4:4
3:3
3:2
2:3
2:2
Passage: 20 25 30 35
2:2 (%): 91 93 95 95
d
UNC1999
Copy Number (Cen:MT)
5:5
4:4
4:3
3:4
3:3
2:2
Passage: 20 25 30 35
2:2 (%): 84 67 50 37
e
UNC1999 + VE821
Copy Number (Cen:MT)
5:5
4:4
3:3
3:2
2:3
2:2
Passage: 20 25 30 35
2:2 (%): 82 75 80 81
Equal MT and centromere
More MT copies
More centromere copies


98

**Figure S5: Related to Figure 5. Homologous Recombination is Required For Inherited MT Amplification**

(a) Treatment of MDA-MB-231 cells with 5 µM VE-821 lowers the level of Chk1 protein by western blot. (n=4 independent treatments). (b) MDA-MB-231 cells treated continuously with 810 nM UNC1999 for 18 passages were then subjected to a combined treatment of 0.81 µM UNC1999 and 5 µM VE-821. (c-e) FISH counts for cells treated with 810 nM of EZH2 inhibitor UNC1999 (c), 5 µM of homologous recombination inhibitor VE-821 (d) or combined treatment (e) shows that inhibiting HR prevents inherited MT amplifications. DNA-FISH data represents 150 cells counted from each passage for each treatment condition.

#### Figure S6

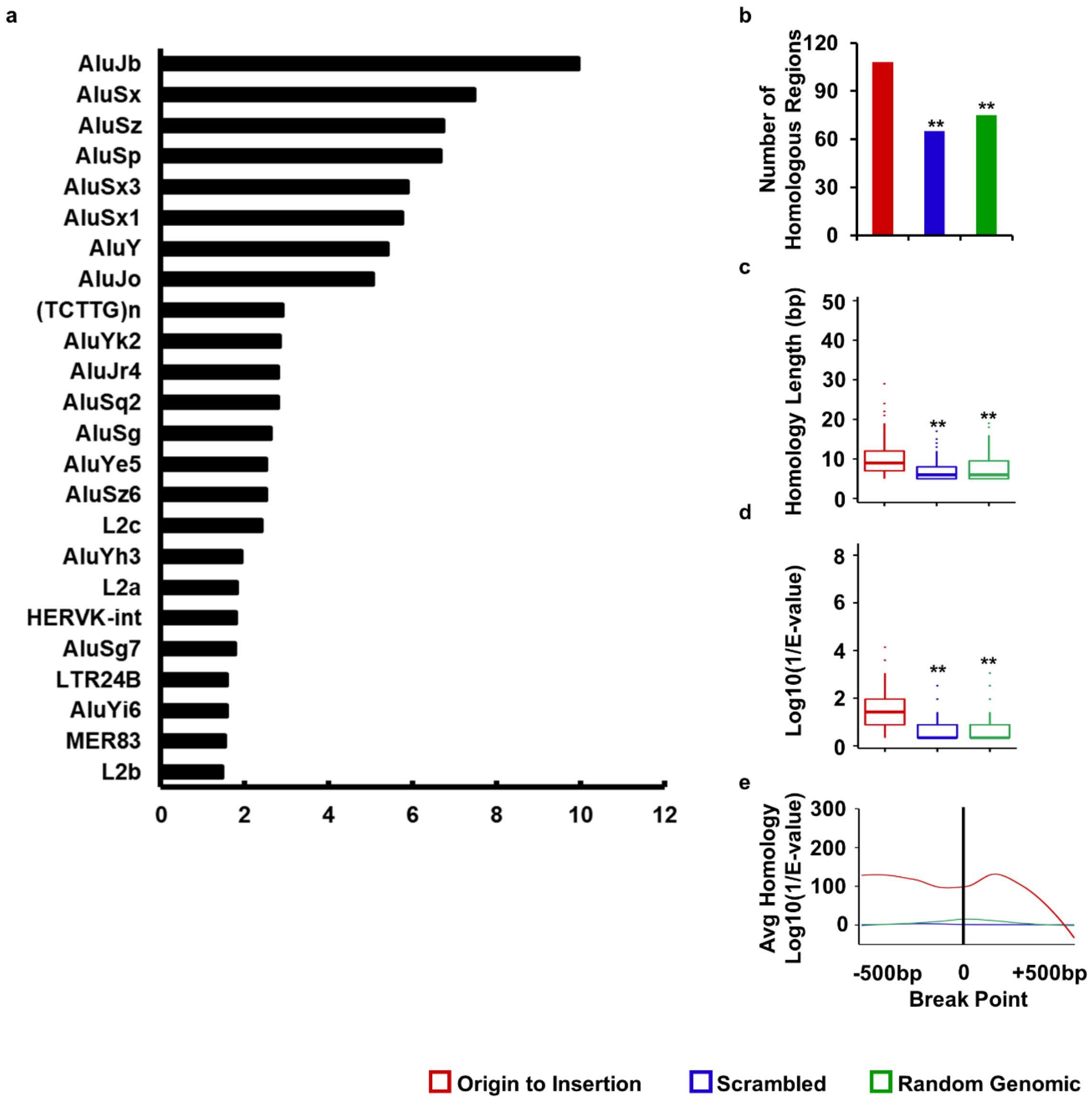
a
AluJb
AluSx
AluSz
AluSp
AluSx3
AluSx1
AluY
AluJo
(TCTTG)n
AluYk2
AluJr4
AluSq2
AluSg
AluYe5
AluSz6
L2c
AluYh3
L2a
HERVK-int
AluSg7
LTR24B
AluYi6
MER83
L2b
0 2 4 6 8 10 12
b
Number of Homologous Regions
0 30 60 90 120
**
**
c
Homology Length (bp)
0 10 20 30 40 50
**
**
d
Log10(1/E-value)
0 2 4 6 8
**
**
e
Avg Homology Log10(1/E-value)
0 100 200 300
-500bp 0 +500bp
Break Point
Origin to Insertion
Scrambled
Random Genomic

**Figure S6: Related to Figure 6. Identification of Interchromosomal Structural Variants with Homology to the MT locus.**

(a) Repeat element enrichment analysis of homologous regions of the MT-insertion-locus reveals significant enrichment of several families of ALU repeat elements are present at the homologous breakpoints. (b) Number of Interchromosomal structural variants with microhomology between the sequence surrounding the break point in the MT locus and the sequence surrounding the break point in the destination locus (Insertion). (c) MT-Insertion pairs have longer microhomologous regions compared to random genomic loci or the scrambled insertion sequence. (d) MT-Insertion pairs have better E-values compared to random genomic loci or the scrambled insertion sequence. (e) The microhomology in MT-insertion pairs is enriched surrounding the GRIDDS predicted break points. ** indicates $p < 1E\text{-}12$ using a Kruskal–Wallis test with a post hoc Wilcoxon–Mann–Whitney test.

### Figure S7

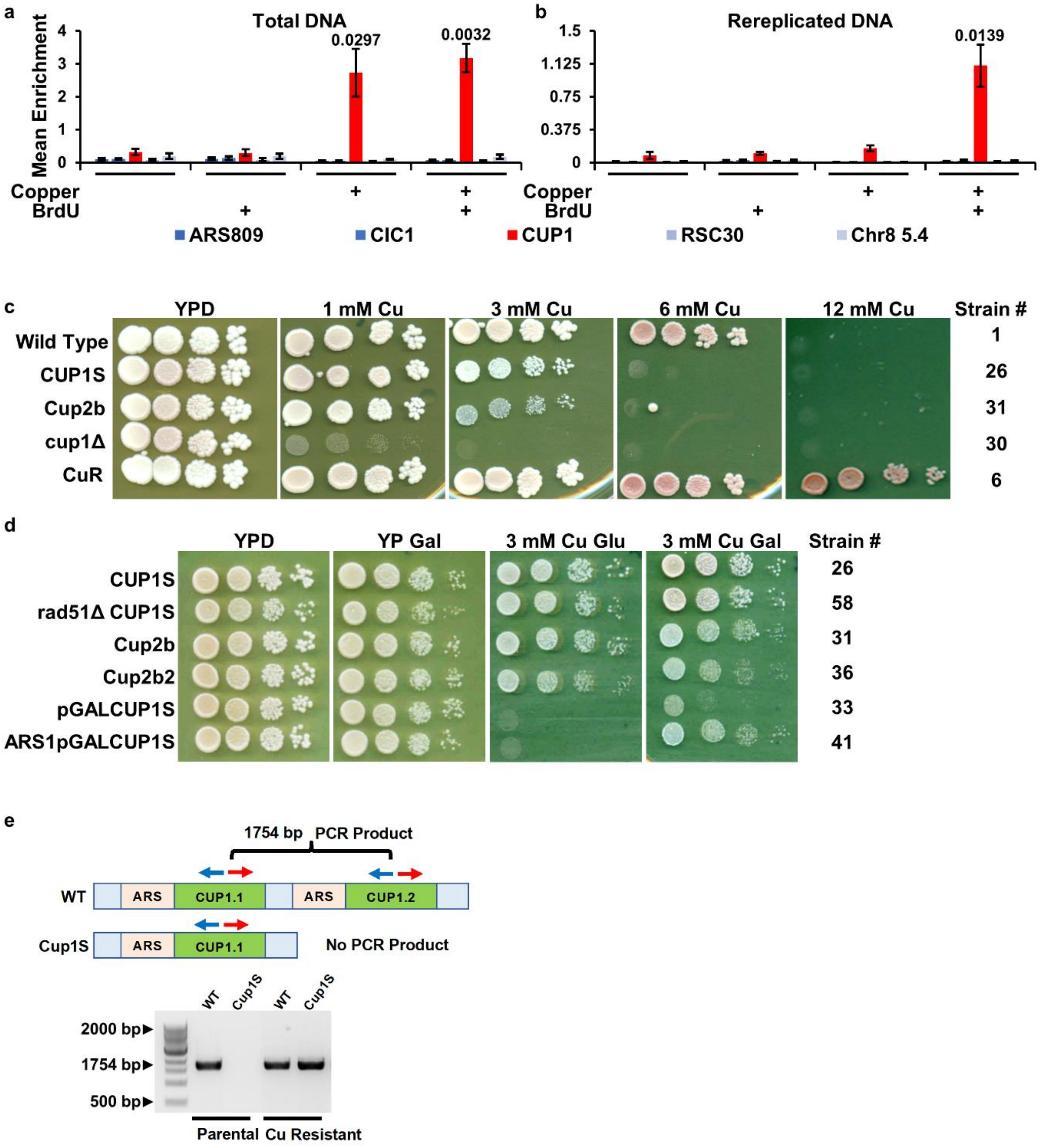


121

#### Figure S7

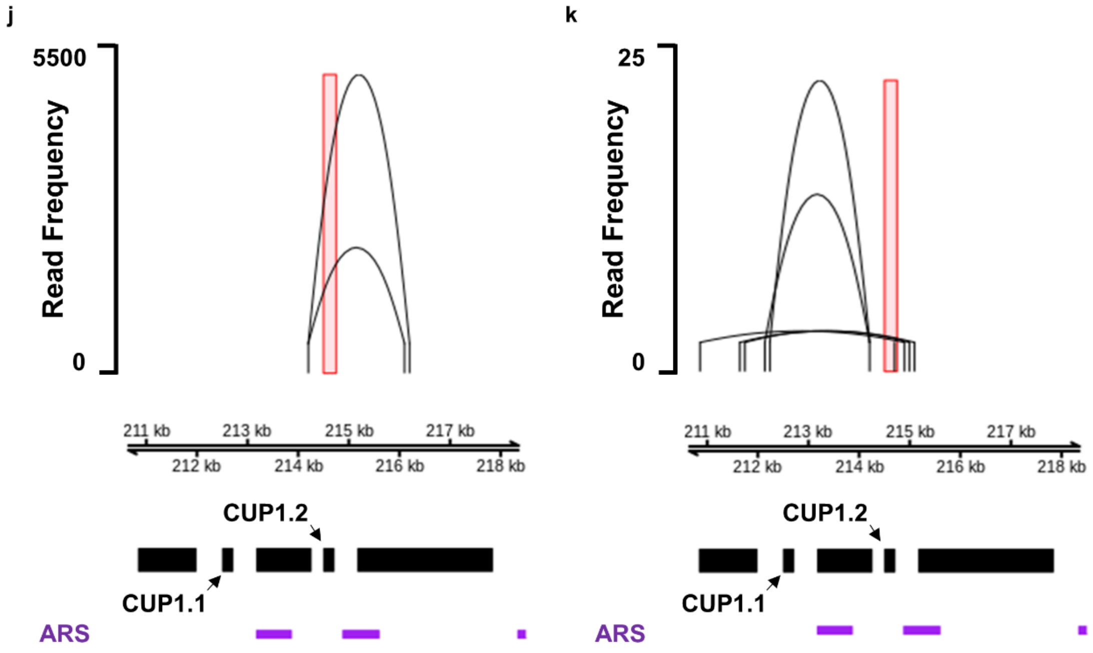


122

**Figure S7**

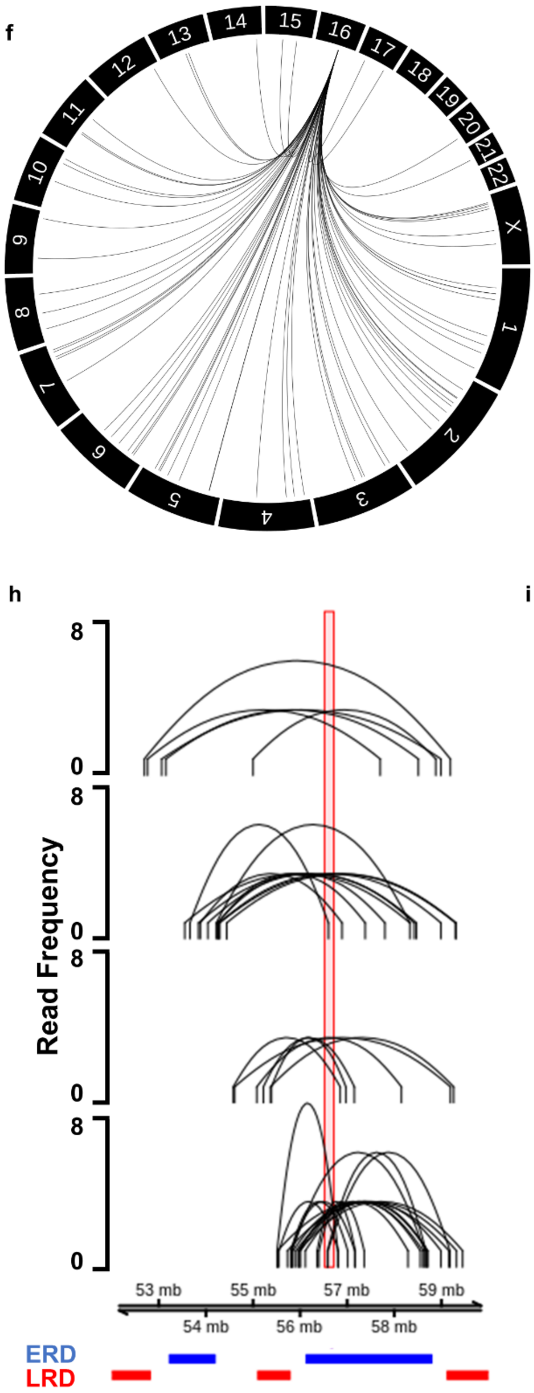


123

**Figure S7: Related to Figure 7. Metal-induced Metallothionein DNA Rereplication is Conserved in Yeast**

(a) Copper induced, amplification of the CUP1 locus. qPCR of input DNA from WT cultures and cultures grown in 6 mM copper for about 30 generations with and without BrdU for Rerep-seq analysis. (n=3 biological replicates) (b) Rereplicated DNA detected at the CUP1 locus. qPCR of Rerep-digested DNA from samples in a. (n=3 biological replicates). qPCR signals were normalized to the mitochondrial gene COX2. Error bars represent the S.E.M. * indicates $p<0.05$ by two-tailed Student's t-test. (c) Yeast spot assay showing sensitivity of mutants to increasing concentrations of copper. Ten-fold serial dilutions of WT (containing at least 2 copies of CUP1), CUP1S (single copy of CUP1), Cup2b (mutation to Cup2 binding motif), cup1Δ (CUP1 deletion), and CuR (copper resistant) on YPD plates with 0, 1, 3, 6, and 12 mM CuSO4. (d) Yeast spot assay demonstrating requirement of CUP1 expression for growth on copper, and functionality of galactose inducible promoter constructs. Ten-fold serial dilutions of CUP1S, Cup2b, Cup2b2, pGALCUP1, and ARS1pGALCUP1, grown on YPD, YP Gal (2% Galactose), YPD + 3 mM CuSO4, or YP Gal + 3 mM CuSO4. (e) Schematic of inverse PCR detection of tandem repeated CUP1. Inverse PCR detecting the reestablishment of a CUP1 tandem amplification in copper adapted CUP1S. (f) Structural variants identified in human Rerep-seq data. (g) Structural variants identified in yeast Rerep-seq data. (h) eccDNA analysis in human cells ranging in size 1-2 Mb. (i) eccDNA analysis in human cells for smaller circles surrounding the MT locus replication domain. (j,k) eccDNA analysis in yeast surrounding the CUP1 locus.

**Table S1** – DNA primers used in this study
